# VINTER: a generative vision–language model for annotating and interrogating spatial tissues

**DOI:** 10.64898/2026.10.01.752251

**Authors:** Jinyu Wu, Wanzhe Xu, Yizhou Zhuang, Yawen Zhang, Martin Loza, Yitao Yang, Kenta Nakai

## Abstract

Spatial transcriptomics links gene expression to tissue structure, yet current models learn embeddings whose meaning is assigned afterwards or read histology without measured expression. Tissue decisions are therefore rarely made from morphology, genes and neighbourhood together, or examined through them. Here we introduce VINTER, a generative vision–language model aligned on 1.45 million histology–caption pairs that reads histology, measured gene expression and spatial neighbourhood as one sequence. It returns the probability of every tissue class, and because genes enter as text, its molecular evidence can be edited with the image held fixed. Reading all three sources together, VINTER annotated tissue more accurately than spatial-domain methods and a spatial foundation model, and transferred across cohorts without target labels. Editing only the neighbourhood resolved the follicular B-cell and plasma-cell programmes behind its reading of tertiary lymphoid structure (TLS) maturation in lung and kidney cancer. Its class probabilities exposed what single labels conceal in triple-negative breast cancer: immune heterogeneity within stroma and cancer hidden beneath benign labels. Without retraining, the same model transferred to Visium HD and single-cell Xenium data and resolved follicular architecture within colorectal cancer TLS. VINTER thus turns spatial annotation into an interrogable reading of tissue across platforms and resolutions.

## Introduction

Spatial transcriptomics measures gene expression within intact tissue, linking molecular states to the histological structures and cellular neighbourhoods that organize disease (1,2). Spatial foundation models and domain-detection methods learn molecular and neighbourhood structure (3–5), image–omics models align histology with transcriptomes (6–8), and generative vision–language models interpret histology through natural language (9,10). These approaches either represent tissue as embeddings or clusters whose meaning is assigned after training, or read images without measured expression. They do not read the morphology of a spot, its genes and those of its neighbours together in one generative decoder, so a tissue decision cannot be both made from all three sources and examined through them.

Two further limitations follow. First, one label per spot flattens the tissue beneath it: stroma assigned to the same class can range from lymphocyte-dense to immune-silent, a difference with prognostic weight in breast cancer (11), and cancer can lie beneath a benign label. Retaining the probability of every class preserves such differences and has improved downstream spatial analyses (12,13). Second, the evidence behind a tissue decision is rarely testable. In silico perturbation of single-cell foundation models nominates genes that shift cell states (14,15), and spatial models have begun to perturb gene pairs (16), but asking which neighbourhood evidence moves a tissue-level judgement requires inputs that separate a spot from its surroundings and can be edited one by one.

Here we introduce VINTER (Vision–language Interrogation of Neighborhood Transcriptomic Evidence and Reasoning), a generative vision–language model that reads histology, measured gene expression and spatial neighbourhood as one sequence. VINTER connects the pathology foundation model UNI (17) to a 7-billion-parameter language decoder aligned on 1.45 million histology–caption pairs from 546 spatial transcriptomic samples. Each spot enters as a haematoxylin and eosin (H&E) image with two ranked lists of gene names, one for the spot and one for its neighbours, and the model decodes its tissue class through dedicated class tokens, returning the probability of every candidate class. Because the molecular evidence enters as text, it stays readable and editable: genes can be removed, demoted or inserted in the spot or its neighbourhood while the image is held fixed.

VINTER annotated cortical layers more accurately than spatial-domain methods and a spatial foundation model, and transferred to an external breast cancer cohort without target labels (18). In lung and kidney cancer, editing only the neighbourhood redirected its reading of tertiary lymphoid structure (TLS) maturation (19), resolving a follicular B-cell programme that raised this reading from a plasma-cell secretory programme that lowered it; in tissue, the two programmes separated the follicular interior of mature TLS from a plasma-cell periphery. In triple-negative breast cancer, the class posterior resolved immune-rich from immune-silent regions within uniformly labelled stroma, revealed T-cell-inflamed and lymphoid-organized environments in genes the model never received, and guided a review that recovered nearly half of the cancer concealed by benign labels. At the 8-µm bin resolution of Visium HD (20), VINTER identified TLS in colorectal cancer accurately and resolved two follicles within a single TLS, with IgG⁺ plasma cells at TLS margins and IgA⁺ plasma cells in the mucosa. VINTER thus turns spatial annotation into an interrogable reading of tissue across platforms and resolutions.

## Methods

### Model architecture and inputs

VINTER combines the histopathology foundation model UNI with the LLaVA-1.5 vision–language model (Vicuna-7B) (17,21). UNI encodes the registered H&E patch centred on each target spot into patch tokens, which a two-layer projector maps into the language embedding space. The text prompt contains the classification request, the candidate classes and, in separately marked spans, the ranked genes of the target spot and of its neighbourhood. Each tissue class is represented by a dedicated token.

For each training fold, the gene panel combines one-versus-rest Wilcoxon markers fitted on the training sections with canonical markers of the tissue. Within the panel, each gene is scored by Eq. (1) on log(1 + counts per 10,000), with μ and σ fitted on training spots, separately for the target spot and for the mean of its six nearest spots. The 20 highest-scoring target genes and the 10 highest-scoring neighbourhood genes enter the prompt as ordered gene-name lists; expression values are not given to the model.

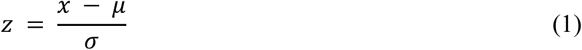

Let **h***_i_* be the final hidden state at the answer position for spot i and **w***_c_* the output row of class token c. The class score, the probability restricted to the task’s candidate classes k and the predicted class are given by Eq. (2), Eq. (3) and Eq. (4). Unless stated otherwise, analyses use these raw probabilities, which express the model’s class support and are not cell-type fractions.

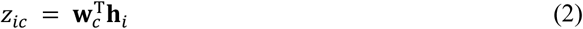

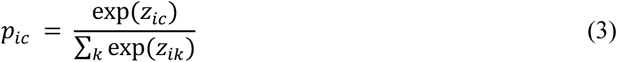

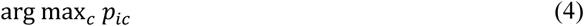

### Model training

VINTER was trained following the two-stage recipe of LLaVA-1.5 (21). The projector was first aligned on 1.45 million HEST histology patches paired with captions generated by Qwen3-VL, excluding all HER2ST and DLPFC samples (22,23); the model was then fine-tuned on the annotation tasks with low-rank adapters (24). The base model was trained on the DLPFC training sections together with labelled HER2ST spots (25), and the TNBC and TLS models were fine-tuned from it. Training used 16 NVIDIA GH200 GPUs.

### DLPFC benchmark

We used the twelve manually annotated DLPFC Visium sections of Maynard et al. (three donors; layers L1–L6 and white matter) (26). Each of the six pairs of adjacent sections was held out in turn (Fig. 2a,b), and the model was trained on six sections (the other pair from the same donor and the four sections of a second donor), excluding the held-out pair from marker selection and normalization. A fixed 35% of held-out spots was used for checkpoint selection and the remaining 30,714 spots were scored. Comparators (BANKSY, spEMO, GraphST, STAGATE, SEDR, STAIG, PROST and SpaGCN (3,7,27–32), and scGPT-spatial with Louvain clustering (33)) were run on each whole section with recommended settings, and their clusters were matched to the reference layers by Hungarian assignment. We report accuracy, macro-F1, balanced accuracy, adjusted Rand index and normalized mutual information as section means. Smoothed VINTER maps reassign a label when a strict majority of its six neighbours agrees on another class. For the held-out pair 151675/151676, image-only and gene-only models were trained in the same way, receiving a neutral filler in place of the genes or a uniform grey image, respectively (Supplementary Fig. 2S3).

**Figure 1.**
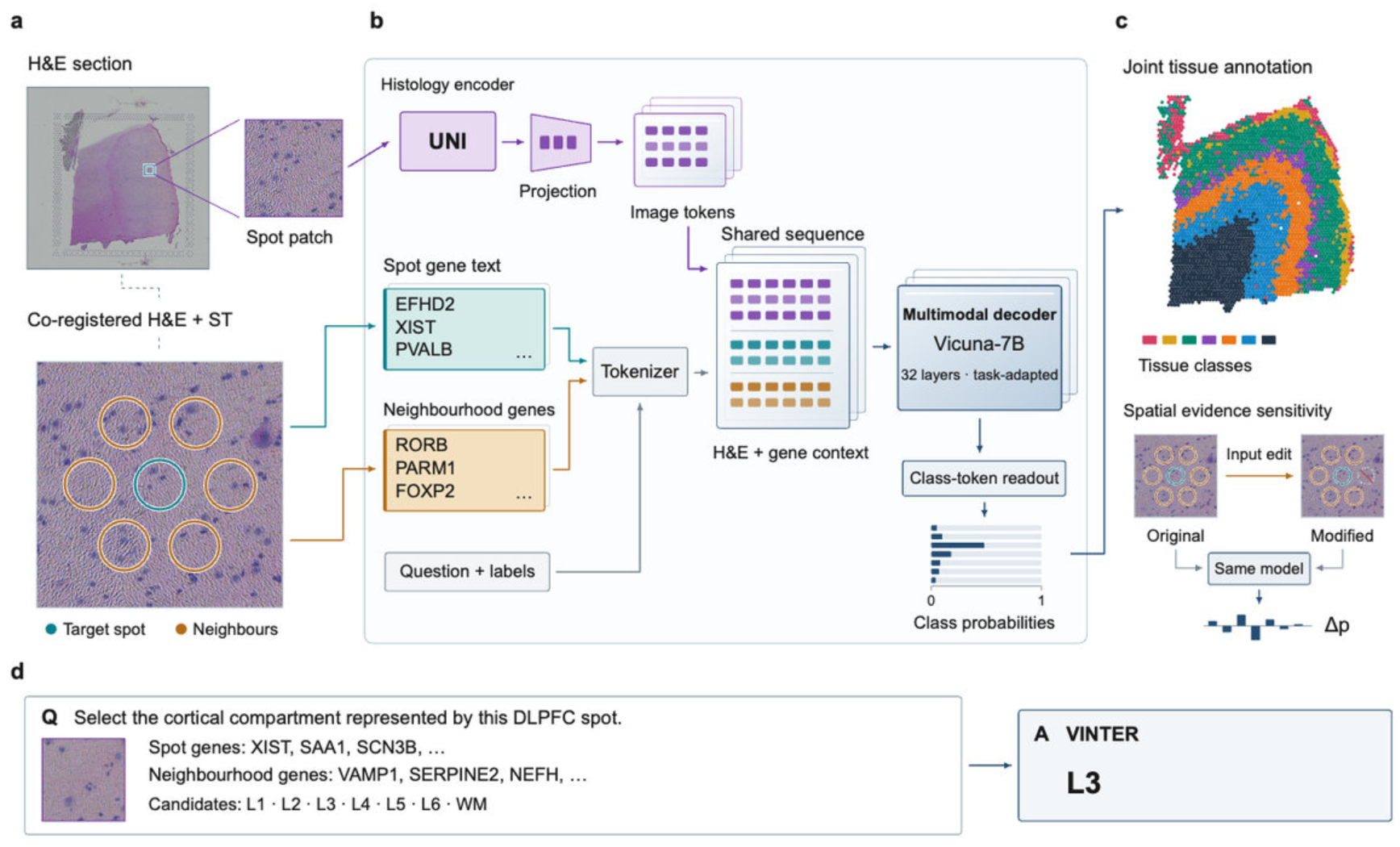
VINTER connects histology and ranked spatial gene evidence in a shared vision–language interface. a, Registered H&E and spatial transcriptomic evidence for a target spot and its six neighbours. The image encoder receives the target-centered patch; the larger neighbourhood crop illustrates spatial registration. b, UNI image tokens are projected into the embedding space of the task-adapted, 32-layer Vicuna-7B decoder and combined with tokenized target-gene and neighbourhood-gene rankings, a classification instruction and candidate labels. The native language-model head provides logits normalized over the predefined class-token set. Representation sheets and the seven probability bars are schematic; the bars describe one spot and sum to one. c, Two uses of the output: a spatial tissue-class map and signed class-probability changes after modification of supplied neighbourhood evidence, as defined in Eq. (5). The map uses raw predictions for 3,563 labelled spots in section 151675; the sensitivity bars are conceptual. Spot-associated class tiles are enlarged for visibility and do not indicate physical capture boundaries. d, An authentic spot from section 151675 illustrates the classification request, abbreviated gene lists, candidate classes and the recorded L3 decision. This panel reconstructs the input presentation and class readout. The a/b image example and d classification example are distinct spots. H&E, haematoxylin and eosin; ST, spatial transcriptomics.

**Figure 2.**
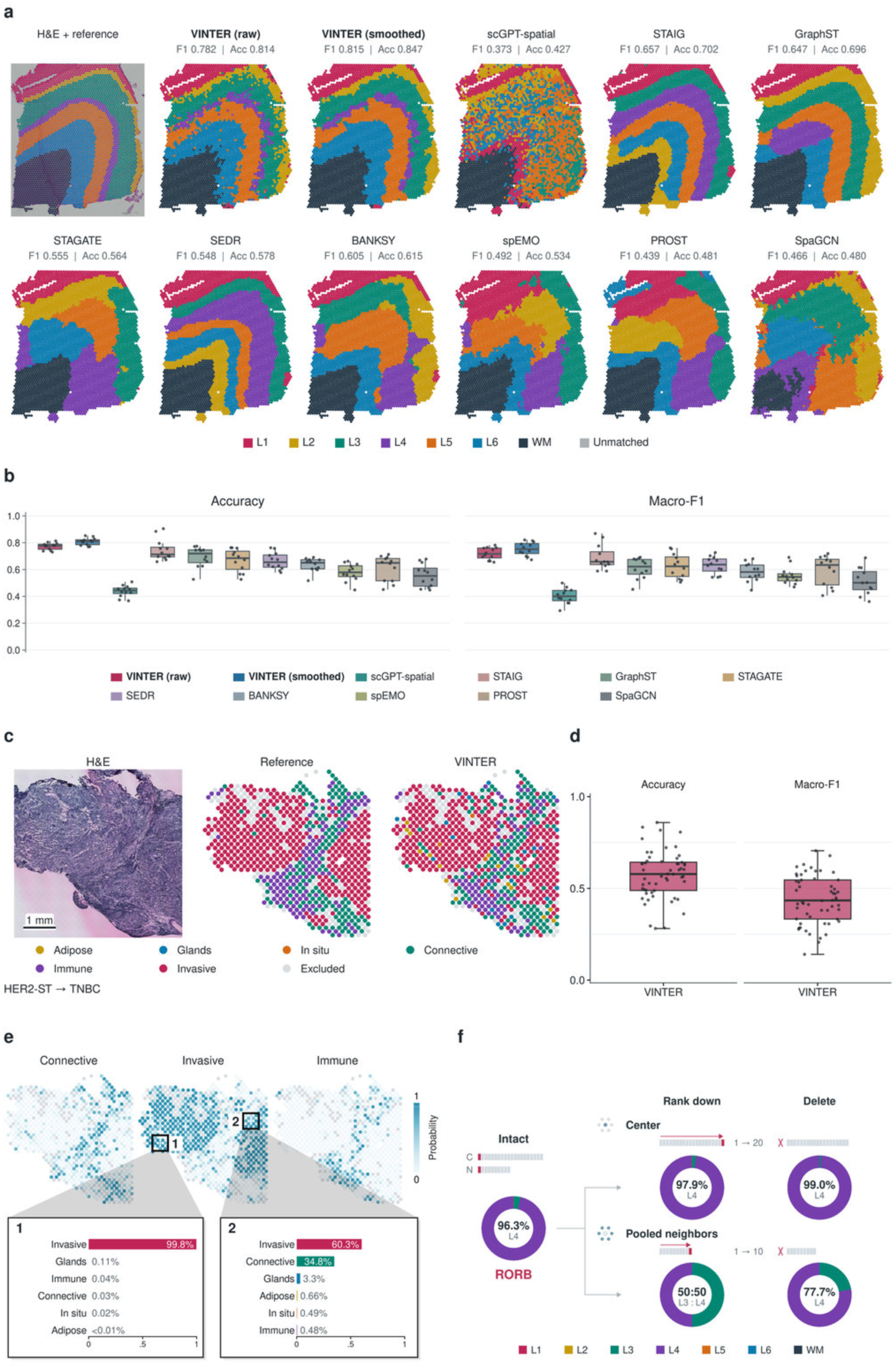
Spatial annotation and functional readouts of VINTER. **a**, Spatial annotation. Reference annotations over H&E and eleven model outputs for DLPFC section 151674. Both raw and six-neighbour-smoothed VINTER outputs are shown. Maps contain all 3,635 annotated spots; attached macro-F1 and accuracy values use the 2,372 canonical EVAL spots. Coordinates, orientation and extent are shared across maps. **b**, Twelve-section benchmark. Accuracy and macro-F1 over 30,714 canonical EVAL spots; balanced accuracy, ARI and NMI are provided in Supplementary Fig. 2S1. One point represents one section; the twelve sections are nested in six held-out section pairs and three donors. Boxes indicate medians and interquartile ranges; whiskers extend to observations within 1.5 interquartile ranges, and all observations are plotted. Macro-F1 averages class F1 over reference classes present in each section; layer-specific F1 is provided in Supplementary Fig. 2S2. There are ten named methods and eleven outputs because VINTER is shown before and after smoothing. **c**, External transfer. A six-class model trained on HER2-ST is applied to TNBC without TNBC labels. Registered H&E, reference annotations and VINTER labels share one coordinate window spanning all measured positions in the illustrative section CN25_E1; 696 of 812 displayed positions are eligible for scoring, and excluded positions are grey. Display disks have a radius of 0.47 times the median nearest-centre distance, without interpolation or new positions. Scale bar, 1 mm. **d**, Patient-level transfer performance. Boxes and points show accuracy and reference-present-class macro-F1 for all 56 test patients (58 sections; 49,574 eligible spots), using the same model as in **c** and pooling each patient’s sections before scoring. Box definitions are as in **b**. **e**, Spatial probabilities and spot-level readouts. Three raw probability channels (Connective, Invasive and Immune) for the section shown in **c** share a 0–1 colour scale, coordinate window and display disks. All 696 eligible spots are coloured; the other 116 positions are grey. The complete six-channel atlas is provided in Supplementary Fig. 2S7. Frames 1 and 2 on the middle map locate two spots (white crosses mark the exact spots read); the linked profiles show all six probabilities of each spot, sorted in descending order, with class colours as in **c**. Both spots have an Invasive argmax: spot 1 has 99.8% Invasive support, whereas spot 2 has 60.3% Invasive and 34.8% Connective support. The spots were selected without reference labels, at the nearest-rank 90th and 10th percentiles of maximum probability within the section’s most abundant predicted class. Supplementary Fig. 2S8 gives the full numeric vectors and the cohort-wide accuracy–retention curve. These raw restricted-softmax probabilities are not calibrated target-domain confidence or cell-type fractions. **f**, Location- and operation-specific input perturbation. RORB is first in both the centre top-twenty and pooled-neighbour top-ten lists of the illustrated L4 spot from DLPFC section 151676. An intact-input reference and a 2 × 2 grid compare the full seven-class probability distributions. Rows specify the edited input block (centre or pooled neighbours); columns specify the operation (move RORB to the final rank or delete its name). Rank-token glyphs indicate the actual edits, with RORB in red; C and N label the intact centre and pooled-neighbour lists. Each edit is applied independently to the intact input; the other input block, image and model are held fixed. Each donut has the same seven-class order and start angle, and each arc occupies its class’s probability without renormalization. Central readouts show L4 support; the neighbour rank edit produces an L3/L4 tie of approximately 49.978% each. The illustrative spot was chosen at the nearest-rank 75th response percentile among fifteen eligible, initially correct L4 spots containing RORB in both lists. These are edits to model inputs, not biological knockouts. The full probability table and the whole-section summary across all 2,214 EVAL spots and 132,840 measured edits are provided in Supplementary Fig. 2S6; the expression-zero-and-rerank sensitivity distribution is provided in Supplementary Fig. 2S4. Multimodal complementarity and target-label adaptation appear in Supplementary Figs. 2S3 and 2S5. VINTER uses supervised whole-pair holdout with TUNE-selected checkpoints, not donor holdout. Clustering baselines are target-section transductive, with annotation-assisted one-to-one Hungarian alignment fitted on the complete annotated section, including EVAL. BANKSY boxes use the per-metric median over eligible resolution runs; maps and layer-specific SI use the fixed displayed run. scGPT-spatial uses a frozen pretrained encoder plus Louvain clustering. TNBC and DLPFC use different ontologies and evaluation units.

### Class probabilities and input edits

Input edits change only the gene-name lists given to the model, with the image and model fixed: a gene is deleted from, or moved to the end of, the centre or the neighbour list (Fig. 2f), or its expression is set to zero before the list is re-ranked (Supplementary Figs. 2S4 and 2S6). The response of each class to an edit is Eq. (5), relative to the intact input. These are edits to the model input, not biological perturbations.

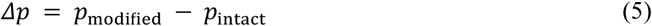

### Transfer to triple-negative breast cancer

The six-class HER2ST vocabulary was transferred to the triple-negative breast cancer (TNBC) cohort of Wang et al. (92 patients; legacy spatial transcriptomics with pathologist pixel annotations) (18) (Fig. 2c,d). A source model fine-tuned on HER2ST training spots was applied without any target labels; TNBC genes were ranked within the HER2ST panel using HER2ST training statistics, and only the target-spot list was supplied. Pathology pixel classes were mapped to the six tissue classes, and spots dominated by one mapped class were scored. Patients were split by a fixed hash into adaptation (18), validation (18) and test (56) sets, and accuracy and macro-F1 were averaged over test patients. Label-budget adaptation used 24–157 labels from adaptation patients (Supplementary Fig. 2S5).

### TLS staging model

We used eight FFPE Visium sections of kidney (KC1–KC3) and lung (LC1–LC5) cancer with author annotations of early TLS, mature TLS, germinal centres (GC), immune infiltrate, tumour and normal tissue (34) (Fig. 3). GC spots were merged into mature TLS, giving five classes (MTLS, ETLS, INFL, TUM and NOR). Because the authors designated KC2 and LC4 as low quality and KC1 contains no mature TLS, maturity was summarized over KC3, LC1, LC2, LC3 and LC5. Each of eight folds held out one section, selected the checkpoint on another section of the same organ and trained on the remaining six. The fold panel combined the training-section markers with 73 curated genes covering germinal-centre B cells, follicular dendritic cells (FDC), T follicular helper (Tfh) cells, T zones, high endothelial venules, B, plasma, T and cytotoxic cells, fibroblastic reticular stroma and antigen presentation.

**Figure 3.**
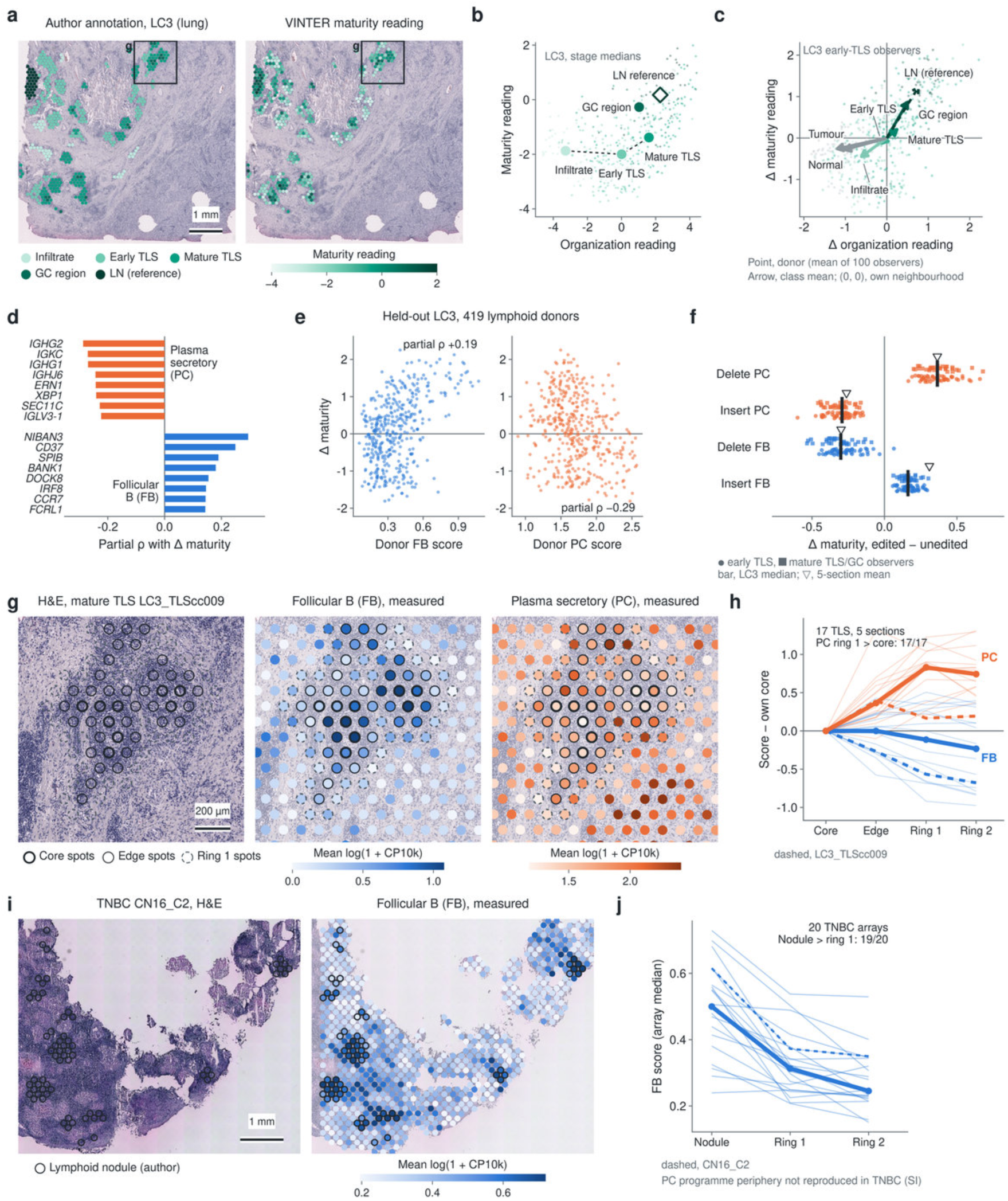
Spatial perturbation with VINTER reveals distinct follicular and plasma-cell programmes around tertiary lymphoid structures. **a**, Representative held-out lung section LC3 (Visium). Left, author annotation of lymphoid spots (infiltrate, early TLS, mature TLS, germinal centre (GC), lymph node (LN)). Right, VINTER maturity reading (logit MTLS − logit ETLS) for TLS and LN spots. Box (g), region shown in **g**. Disc diameter, 0.94 × the centre- to-centre spacing. **b**, LC3 lymphoid spots on the two readouts (small points) and per-stage medians (large points): organization reading, (logit ETLS + logit MTLS)/2 − logit INFL; maturity reading as in **a**. The dashed line joins the infiltrate, early-TLS and mature-TLS medians as a reading guide. GC region, germinal-centre spots within mature TLS. Diamond, lymph node reference; the lymph node was not a training target and is not a TLS stage. **c**, Real-neighbourhood swap. The 100 LC3 early-TLS observers kept their own H&E patch and centre gene list and received the neighbour gene list of one real donor spot from the same section. Axes give the change of both readouts from the observer’s reading with its own neighbourhood (origin, no change). Points, the 500 LC3 donors (each the mean over the 100 observers); arrows, mean per donor class, with donors weighted equally (dashed, LN reference donors). Across the five primary sections (class mean per section, then an equal-weight mean over the sections that have such donors), the maturity shift was −0.46, −0.40 and −0.41 for infiltrate, normal and tumour donors, 0.00 for early-TLS donors, +0.43 for mature-TLS donors, +0.60 for GC donors (2 sections) and +1.22 for LN donors (1 section). **d**, Two programmes extracted from the donor states that move the reading: follicular B (FB) and plasma secretory (PC), top 8 of 20 genes each. Bars, partial Spearman correlation between a lymphoid donor neighbourhood’s measured expression and the Δ maturity it causes, with neighbourhood B-cell, T-cell and *PTPRC* expression partialled out (mean of kidney and lung). **e**, LC3 held out: both programmes re-derived on the other four sections and scored on the 419 LC3 lymphoid donors. Axes show the raw donor programme score and the Δ maturity reading from **c**; partial ρ as in **d**. Across the five held-out sections and three observer–readout combinations, FB was positive (+0.06 to +0.39) and PC negative (−0.24 to −0.49) in 15 of 15 tests. **f**, Bidirectional editing of real donor neighbour lists. Programme genes present in the list were deleted, or the donor’s three highest-ranked programme genes were inserted at, or promoted to, the top of the list (insertions keep the list length; deletions shorten it). Δ maturity is relative to the same donor unedited. Each point is one LC3 observer’s mean effect over its donors (circles, 50 early-TLS observers; squares, 30 mature-TLS/GC observers: 26 MTLS, 4 GC). Bar, median of the 80 LC3 observers. Triangle, mean over five sections and both observer groups, each weighted equally. In total, 351 observers and 34,697 observer– donor pairs. Every edit shifted the reading in the expected direction in all five sections; the share of observers shifted in that direction, computed for each edit within each observer group and averaged over sections, was 95–100%. Deletions with no programme gene in the list are excluded. **g**, Mature TLS LC3_TLScc009: H&E and measured FB and PC programme scores (mean log(1 + CP10k) over programme genes). Thick ring, TLS core spots (all six neighbours in the same TLS); thin ring, TLS edge spots; dashed ring, first ring of non-TLS spots. Discs at the true 55-µm spot size. Colour scales span the 5th–97th percentile of the spots whose centres lie in the panel and are set independently per panel. **h**, Programme score minus the object’s own core score in the 17 mature TLS of the five primary sections that have a core, an edge and two outer rings. For each TLS and compartment, the spot median is used. Thin lines, individual TLS; thick lines, medians of these paired differences; dashed, LC3_TLScc009. PC is higher in ring 1 than in the core in 17 of 17 TLS; FB is higher at the edge than in ring 1 in 11 of 17. **i**, Independent TNBC cohort of 92 primary tumours profiled by legacy ST (18), array CN16_C2 (the array with the most nodule spots): registered H&E and measured FB score (colour scale as in **g**, set for this panel). Rings mark nodule spots: at least half of the spot is tissue, at least half of that tissue is author-annotated lymphoid nodule, and the spot has counts. **j**, FB score (array median) in nodule spots and in the first and second rings around them, for the 20 primary arrays with nodules. Thick line, median; dashed, CN16_C2. FB is higher in nodules than in ring 1 in 19 of 20 arrays, and higher in nodules than in ordinary immune infiltrate in 17 of the 18 patients with both. The FB programme shares 5 of 20 genes with the authors’ own TLS signature (*BLK*, *CCR7*, *CD37*, *FCMR*, *NIBAN3*; hypergeometric P = 9.9 × 10⁻¹¹; 28 signature genes in a 17,845-gene universe). The PC programme periphery was not reproduced in this cohort: the PC score was higher in ring 1 than in nodules in 6 of 20 arrays (Supplementary Fig. 3S6). Scale bars: 1 mm (**a**, **i**), 200 µm (**g**). Visium rings are hexagonal hops from the TLS. On the legacy-ST checkerboard, ring 1 is the diagonal neighbours and ring 2 the next axial layer (±2, 0)/(0, ±2). Perturbations edit the gene-name lists given to the model and read its judgement; they are not tissue interventions.

Two readouts were computed from the class scores of Eq. (2): organization, Eq. (6), and maturity, Eq. (7). Organization was evaluated per spot (TLS versus infiltrate). For maturity, TLS objects were connected regions of TLS-labelled spots, called mature when they contained an MTLS or GC spot, and AUCs were averaged over sections. A variant was trained without 21 germinal-centre B, FDC and Tfh genes (Supplementary Fig. 3S1).

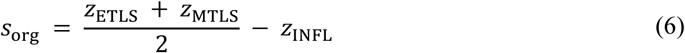

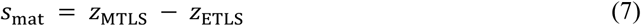

### Neighbourhood perturbation and programme discovery

Perturbations used the five-class model and changed only the gene lists. Early-TLS, infiltrate and mature-TLS spots of each section served as observers, and lymphoid spots, topped up with tumour and normal spots, as donors. In a real-neighbourhood swap, an observer kept its H&E patch and centre list and received a donor’s neighbour list from the same section (Fig. 3c); single-gene deletions and demotions and programme-level deletions, demotions and insertions were scanned in the same way (Supplementary Fig. 3S2).

To identify the neighbourhood states that move the reading (Fig. 3d), each lymphoid donor’s maturity response was averaged over the early-TLS observers of its section and correlated within sections with the measured whole-transcriptome expression of the donor’s six neighbours, each gene scored by Eq. (8), adjusting for neighbourhood B-cell, T-cell and PTPRC expression by partial correlation. Genes with consistent signs in kidney and lung and in at least four of five sections were clustered by co-expression, and the leading coherent cluster in each direction was trimmed to 20 genes, giving the follicular B (FB) and plasma secretory (PC) programmes. Leave-one-section-out validation re-derived the programmes from four sections and tested them on the fifth (Fig. 3e). In bidirectional editing (Fig. 3f), programme genes were deleted from a donor’s neighbour list, or the donor’s three highest-ranked programme genes were inserted at or promoted to the top of the list.

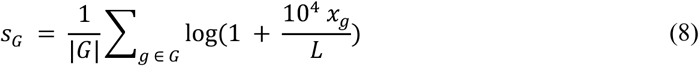

### Spatial localization and replication in TNBC

Within mature TLS objects, core spots had all six neighbours in the object, the other TLS spots formed the edge, and the first and second layers of surrounding spots formed rings 1 and 2 (Fig. 3g,h). In the TNBC cohort, FB and PC were scored on whole-transcriptome counts of primary specimens (Fig. 3i,j); nodule spots were spots whose tissue was mostly annotated as lymphoid nodule, and rings were taken on the legacy spatial transcriptomics lattice. Overlap with the authors’ TLS signature (18) was tested with a hypergeometric test.

### Class posteriors in TNBC

Posterior analyses used the zero-shot source model and the 92 primary specimens (Fig. 4). Programmes were scored by Eq. (8) on measured whole-transcriptome counts, excluding the 80 model-input genes unless stated. Within spots labelled connective tissue, p(immune) was compared with the pathologists’ high- and low-TIL stroma annotation by within-patient AUC and Spearman correlation. Stromal spots were split into within-patient tertiles of p(immune), and hot and cold pseudobulks were compared with a paired edgeR quasi-likelihood model (35) and by published programmes (antigen processing, IFN-γ-related genes, cytolytic activity, lymphoid-organizing chemokines and fibroblast markers) (36–40). Hot stroma within two spot spacings of a lymphoid nodule was compared with hot stroma more than four spacings away, excluding spots containing nodule pixels.

**Figure 4.**
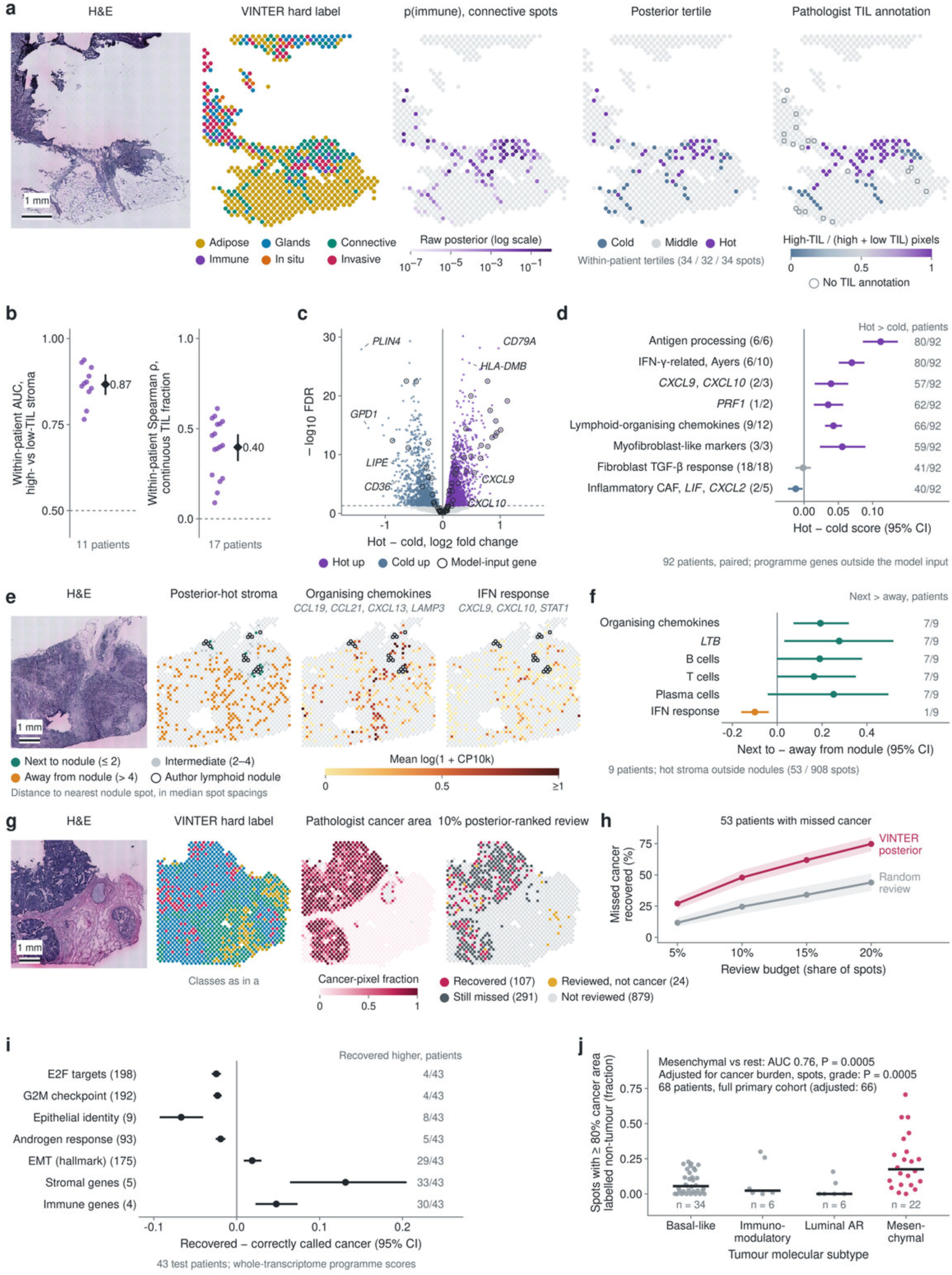
VINTER’s class probabilities reveal stromal immune heterogeneity and recover overlooked cancer in triple-negative breast cancer. VINTER trained on HER2ST was applied without TNBC training to primary tumours from 92 patients profiled by legacy spatial transcriptomics with pixel-level pathology annotation (18). Raw (uncalibrated) class posteriors are shown throughout. Programme and gene readouts come from the measured whole transcriptome. Programmes in d and genes labelled in c exclude the 80 model-input genes; readouts in f and i are scored on all measured members, including model-input genes where present. **a**, Representative tumour (patient 4, section CN2_D2). From left: H&E; VINTER hard labels for all 629 spots; raw p(immune) for the 100 analysed spots labelled connective tissue (log colour scale; other spots grey); within-patient tertiles of p(immune); pathologist annotation, shown as the high-TIL share of high-plus low-TIL stroma pixels (open circles, connective spots without TIL annotation). Case chosen for coverage and balance of high- and low-TIL annotation, not for agreement. Scale bar, 1 mm (from the nominal array pitch: two array-x steps ≈ 200 µm). **b**, Agreement between p(immune) and pathologist TIL annotation within connective-labelled stroma. Left, within-patient AUC for spots with a majority of high- versus low-TIL pixels (11 patients with at least five spots of each). Right, within-patient Spearman correlation with the continuous high-TIL share (17 evaluable patients). Dots, patients; diamonds and bars, patient mean and 95% patient-bootstrap interval (10,000 resamples). **c**,**d**, Posterior-hot versus cold stroma (top versus bottom within-patient tertile of p(immune)), paired whole-transcriptome pseudobulk in 92 patients. **c**, Paired edgeR quasi-likelihood test (12,667 genes; 2,059 higher in hot and 1,261 higher in cold stroma at FDR < 0.05); hot up and cold up, FDR < 0.05; open circles, genes in the model input panel. **d**, Programme scores (mean log(1 + CP10k) over members measured in every sample); in parentheses, measured / prespecified members after exclusion of model-input genes: antigen processing (36), IFN-γ-related (37; *IFNG*, *CXCL11*, *IDO1* and *GZMA* not measured), *CXCL9*/*CXCL10* (*CXCL11* not measured), *PRF1* (cytolytic activity; *GZMA* not measured), lymphoid-organising chemokines (39; 12-chemokine set; *CCL18*, *CCL21* and *CXCL11* not measured), myofibroblast-like markers, fibroblast TGF-β response (Pan-F-TBRS; the model-input gene *TAGLN* removed from its 19 genes) and inflammatory CAF markers (*LIF*, *CXCL2*; *IL6*, *IL11* and *CXCL1* not measured). Points and bars, mean hot − cold difference and 95% patient-bootstrap interval, coloured when the interval excludes zero; numbers, patients with a higher score in hot stroma. **e**,**f**, Posterior-hot stroma next to and away from author-annotated lymphoid nodules. **e**, Patient 16 (section CN8_E2; the patient with the most nodule-free hot stromal spots next to a nodule): H&E; hot stromal spots that contain no nodule pixel, coloured by distance to the nearest nodule spot in median nearest-neighbour spot spacings (next to, ≤ 2; intermediate, 2–4, shown but not compared; away, > 4); mean expression of lymphoid-organising chemokines (*CCL19*, *CCL21*, *CXCL13*, *LAMP3*) and of IFN-response genes (*CXCL9*, *CXCL10*, *STAT1*) in the same spots. Open circles, nodule spots (lymphoid-nodule pixels > half of the spot foreground). **f**, Next-to minus away difference in nine patients with at least three hot stromal spots in each group (53 and 908 spots). Readouts: organising chemokines as in e; *LTB*; B cells (*MS4A1*, *CD79A*); T cells (*CD3E*, *CD3D*, *CD2*, *TRAC*); plasma cells (*JCHAIN*, *MZB1*, both model-input genes); IFN response as in e. Scores are spot-level mean log(1 + CP10k), averaged within patient and zone; bars, 95% patient-bootstrap interval; numbers, patients with a higher score next to nodules. **g**,**h**, Posterior-guided review of cancer concealed by benign hard labels. **g**, Patient 96 (section CN48_E1): H&E; VINTER hard labels (colours as in a); pathologist cancer-pixel fraction (tumour, tumour region and in situ pixels over spot foreground); outcome of reviewing 10% of the patient’s spots, chosen among spots labelled non-tumour in order of p(invasive) + p(in situ). Recovered and still missed, spots with a majority of cancer pixels that the hard label called non-tumour; not reviewed, other valid spots; background grey, spots outside the review-eligible set. **h**, Mean recovery of such missed cancer across 53 test patients with at least one miss, for posterior-ranked review and a fixed random order at the same budget (share of each patient’s valid spots); bands, 95% patient-bootstrap interval. **i**, Programme scores of cancer spots recovered by the 10% review minus cancer spots the hard label called correctly, in 43 test patients with at least three spots of each (paired pseudobulk; Hallmark E2F targets, G2M checkpoint, androgen response and epithelial–mesenchymal transition; epithelial identity, *CDH1*, *EPCAM*, *KRT8*, *KRT18*, *KRT19*, *CLDN4*, *CLDN7*, *ESRP1*, *GRHL2*; stromal genes, *COL1A1*, *COL1A2*, *COL3A1*, *DCN*, *LUM*; immune genes, *PTPRC*, *CD3E*, *CD2*, *CD68*; measured members in parentheses; G2M, androgen response, EMT, epithelial identity and stromal genes each include measured model-input genes). Recovered and correctly called spots had a similar pathologist cancer area (patient means 0.71 and 0.74). Bars, 95% patient-bootstrap interval; numbers, patients with a higher score in recovered cancer. **j**, Fraction of spots with ≥ 80% cancer pixels that the hard label called non-tumour, by the authors’ tumour molecular subtype, in 68 patients with at least ten such spots (full primary cohort, not restricted to the test patients in h and i); bars, medians. Mesenchymal versus other subtypes: AUC 0.76 (Mann–Whitney *P* = 0.0005); rank regression adjusted for whole-section cancer area, log number of cancer spots and grade, *P* = 0.0005 (HC3 errors; 66 patients with complete covariates). Spots are tissue mixtures; programme scores are tissue-level relative abundances, not cell-intrinsic activities. Scores in d and i are computed on pseudobulk, those in f on spots, so effect sizes are not compared across panels.

Missed cancer was defined as spots with a majority of cancer pixels but a non-tumour hard label. Within each patient, non-tumour spots were reviewed in order of p(invasive) + p(in situ) up to a budget expressed as a fraction of the patient’s spots, and recovery was compared with random review. Recovered and correctly called cancer were compared by paired pseudobulk programme scores (including Hallmark sets (41)). The fraction of high-cancer spots called non-tumour was compared across the authors’ molecular subtypes (18) with the Mann–Whitney test and with rank regression adjusted for cancer area, cancer spot number and grade.

### Zero-shot application to Visium HD and Xenium

We used colorectal cancer patients P1, P2 and P5 (20). For each patient, Visium HD and Xenium were performed on adjacent sections of the same FFPE block. For Visium HD, we used the authors’ RCTD cell-type annotation (42) and their tumour periphery labels. The eight fold models of the TLS staging model were applied without retraining, and their logits were averaged (Fig. 5). The readouts were maturity (Eq. (7)), organization (Eq. (6)) and the TLS probability, the sum of the MTLS and ETLS probabilities.

**Figure 5.**
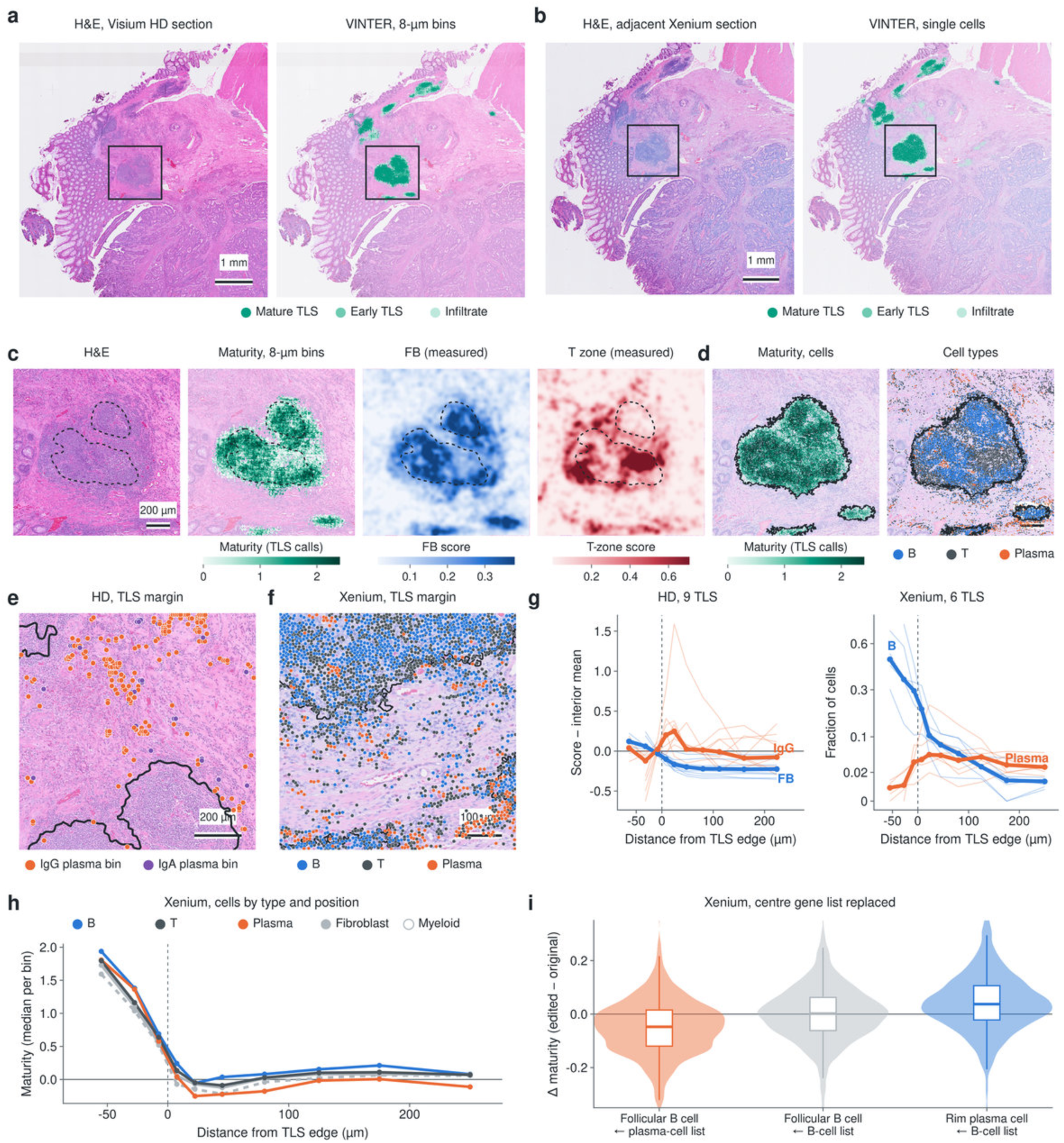
VINTER reads TLS state on colorectal cancer Visium HD bins and Xenium cells without retraining. The TLS model of Fig. 3, trained on lung and kidney cancer Visium, was applied without retraining to colorectal cancer patient P1 (20). This patient was profiled by Visium HD and, on an adjacent section, by Xenium. Each 8-µm bin or segmented cell was read with its real spatial neighbours and the surrounding H&E (Methods). **a**, Visium HD section: H&E and VINTER lymphoid calls on 8-µm bins. **b**, Adjacent Xenium section: H&E and VINTER lymphoid calls on single cells. **c**, Largest TLS on HD: H&E, maturity on TLS-called bins, and the measured follicular B (FB) and T-zone programmes. Dashed lines outline follicles defined by the measured FDC programme. **d**, Largest TLS on Xenium: maturity on TLS-called cells, and B, T and plasma cells. Lines outline VINTER TLS objects. **e**, TLS margin on HD: author-annotated plasma bins coloured by dominant isotype (IgG or IgA). **f**, TLS margin on Xenium: B, T and plasma cells. **g**, Profiles by distance from the TLS edge for HD (left) and Xenium (right) TLS objects. Thin lines are individual objects; thick lines are medians. **h**, Xenium: maturity of each cell type by distance from the TLS edge. **i**, Change in maturity when only the centre gene list of a Xenium cell is replaced. Follicular B cells received a plasma cell’s list or another B cell’s list; rim plasma cells received a B cell’s list. Medians with 95% CIs are shown.

Inputs followed training. Gene lists were ranked within each fold’s panel using that fold’s frozen training statistics, and the image was the 150-µm H&E field around the unit. On Visium HD, centre units were in-tissue 8-µm bins, and the neighbourhood was the eight surrounding bins. On Xenium, centre units were segmented cells with sufficient counts, and the neighbourhood was the six nearest cells; cell positions were mapped to the H&E image with the 10x image alignment. The Xenium panel covers only part of each fold’s gene panel; absent genes entered the ranking with zero counts.

Xenium cells were typed by panel marker genes (Supplementary Table 5S1). TLS objects were connected regions of TLS-called bins (Visium HD) or cells (Xenium) above a minimum size, and distances were signed distances to the object edge. Measured programme scores were the mean over genes of log1p CP10k (Eq. (8)) and were spatially smoothed for display and for the correlation with maturity.

On Visium HD, the authors’ RCTD plasma bins were typed as IgG- or IgA-dominant by their own immunoglobulin heavy-chain counts. Their density was expressed per stromal bin, by distance from TLS. Compartments (TLS and rim, tumour border and mucosa) were assigned from TLS calls, the authors’ periphery labels and the local epithelial content.

For cross-platform registration in P1, the Xenium H&E image was registered to the Visium HD H&E image by a similarity transform fitted to matched image keypoints. TLS calls were then compared between each Xenium cell and the nearest Visium HD bin, and each Xenium TLS object was matched to the Visium HD object that contains most of its cells.

For centre-gene-list replacement in Xenium P1, only the centre gene list of follicular B cells and of plasma cells at the TLS margin was replaced by that of a randomly drawn donor cell, taken from the same fold. The H&E field and the neighbourhood list were left unchanged. Changes are edited minus unedited, paired by cell, and are reported as medians with bootstrap 95% CIs.

### Statistics and software

Sections and patients were the units of analysis for comparisons between tissue states; spots nested within them were not treated as independent replicates. Unless stated otherwise, confidence intervals are percentile intervals from 10,000 patient-level bootstrap resamples; for centre-list swaps, which compare each target with itself, intervals were bootstrapped over targets. Analyses used Python with PyTorch, Transformers, PEFT, Scanpy (43), Squidpy (44), scikit-learn and SciPy, and R with edgeR and ggplot2.

## Results

### A generative vision–language interface connects histology with local molecular evidence

Spatial tissue annotation requires an interpretation of morphology in its molecular context. We developed VINTER (Vision–language Interrogation of Neighborhood Transcriptomic Evidence and Reasoning) to connect these two forms of evidence through a shared vision–language sequence, adapting a generative vision–language backbone to spatially resolved tissue classification (Fig. 1a,b). For each target spot, the model receives a registered haematoxylin and eosin (H&E) patch, an ordered list of genes from the target spot and a separate list summarizing its six spatial neighbours. A natural-language classification request and an explicit map between candidate tokens and tissue labels specify the task. This interface places image features and human-readable molecular evidence in a common sequence while preserving their target and neighbourhood origins.

VINTER combines the histopathology foundation model UNI with the 32-layer Vicuna-7B decoder of LLaVA-1.5 (17,21). The image encoder supplies 196 patch tokens, which a trainable projection maps into the language-model embedding space. In parallel, gene symbols are ordered using training-fitted expression scores and tokenized as text: the cortical task uses 20 target genes and 10 neighbourhood genes. The resulting sequence is processed by a decoder adapted with low-rank updates. Pathology-image alignment through projector training establishes the visual interface, followed by optimization for tissue-class decisions. Morphology, molecular identity and local spatial context thus converge in the decoder and are read out through its native language-model head.

The native language-model head provides both the tissue assignment and the complete vector of candidate-class probabilities. For cortical annotation, seven atomic output tokens correspond to layers L1–L6 and white matter (WM); their logits are normalized within this candidate set. These outputs connect spatial annotation to evidence interrogation: the most-supported class defines the tissue map, while the full probability vector records how class support is redistributed when the supplied evidence changes (Fig. 1c). An authentic spot example illustrates the transition from an image and ranked gene lists to an L3 assignment (Fig. 1d). We used this interface to evaluate multimodal annotation, identify the molecular and spatial inputs supporting tissue decisions, and extend the resulting probabilities into spatial readouts.

### VINTER provides accurate and transferable tissue annotation with readable probabilities and editable inputs

We first benchmarked VINTER on the laminar organization of the human dorsolateral prefrontal cortex (DLPFC), using the manually annotated spatial transcriptomics sections of Maynard et al. (26). Twelve sections from three donors were evaluated as six held-out section pairs: VINTER was trained on sections outside each pair, a fixed 35% TUNE partition of the held-out pair was reserved for checkpoint selection, and the remaining 30,714 canonical EVAL spots were scored. We compared VINTER with eight spatial-domain methods (3,7,27–32) and the foundation model scGPT-spatial, whose clusters were matched to the reference layers section by section, and report raw and six-neighbour-smoothed VINTER outputs separately (Fig. 2a,b).

Across the twelve sections, raw VINTER achieved a mean macro-F1 of 0.719 and accuracy of 0.773, and smoothing raised them to 0.753 and 0.808. The strongest comparator, STAIG, reached a mean macro-F1 of 0.690 and accuracy of 0.745 (Fig. 2b). In the representative section 151674, VINTER recovered the six cortical layers and white matter (WM) with a macro-F1 of 0.782 (0.815 after smoothing; Fig. 2a). Balanced accuracy, adjusted Rand index, normalized mutual information and layer-specific F1 are provided in Supplementary Figs. 2S1 and 2S2. Separately trained image-only and gene-only models showed that the two inputs are complementary: on 4,541 EVAL spots, the joint model was correct for 77.93% of spots, compared with 75.47% for the gene-only and 65.69% for the image-only model, and it correctly assigned 130 of the 620 spots that both single-modality models misclassified (Supplementary Fig. 2S3).

We then asked whether a tissue vocabulary learned in one cohort transfers to another without target labels. We applied the model trained on HER2-positive breast cancer (25) to 92 patients with triple-negative breast cancer (TNBC) profiled with paired H&E and spatial transcriptomics (18), retaining the same six tissue classes. In the 56 test patients (58 sections; 49,574 eligible spots), VINTER achieved a patient-mean macro-F1 of 0.438 (95% patient-bootstrap interval, 0.404–0.472) and accuracy of 0.574 (0.539–0.607) without any TNBC labels (Fig. 2d). In section CN25_E1, the predicted invasive cancer, connective tissue and immune infiltrate reproduced the organization of the reference annotation (section macro-F1, 0.678; Fig. 2c). With 157 target labels, LoRA adaptation raised the patient-mean macro-F1 to 0.484 (Supplementary Fig. 2S5).

Beyond the winning label, the native language-model head returns the full probability vector over the candidate classes (Fig. 2e). In CN25_E1, the invasive-cancer, connective-tissue and immune channels outlined the tumour, its stroma and the immune compartment. Two spots with the same invasive-cancer label differed in their alternatives: one assigned 99.8% of its support to invasive cancer, whereas the other assigned 60.3% to invasive cancer and 34.8% to connective tissue. Across the 56 test patients, retaining spots whose maximum probability exceeded 0.8 kept 61.8% of spots at an accuracy of 65.0%, compared with 57.2% for all eligible spots (Supplementary Figs. 2S7 and 2S8). We return to these probabilities in Fig. 4 to read tissue states that a single label conceals.

Because molecular evidence enters VINTER as ordered gene names, each input can be edited and the resulting change in class support read directly (Fig. 2f). In an L4 spot of DLPFC section 151676 in which RORB ranked first in both the centre and the pooled-neighbour list, the intact input assigned 96.3% of its support to L4. Moving RORB to the end of the centre list or deleting it left L4 support at 97.9% and 99.0%, whereas the same edits in the neighbour list reduced it to an L3/L4 tie (50.0% each) and to 77.7%. Across all 2,214 EVAL spots and 132,840 single-gene edits of this section, neighbour deletions produced the largest median change in class support (Supplementary Figs. 2S4 and 2S6). The model’s judgement therefore depends on where a gene appears in its evidence as well as on which gene it is, a property we use in Fig. 3 to dissect a tissue decision.

### Spatial perturbation with VINTER reveals distinct follicular and plasma-cell programmes around tertiary lymphoid structures

Tertiary lymphoid structures (TLS) mature from lymphoid aggregates into follicles with germinal centres, and mature TLS generate antibody-producing plasma cells (45). Because VINTER receives the neighbourhood as an editable gene list, we used it to ask which neighbourhood states the model uses to read TLS maturation, and where those states lie in tissue. We trained a five-class model (mature TLS including germinal centres, early TLS, immune infiltrate, tumour and normal tissue) on eight lung and kidney cancer Visium sections with author TLS annotations (34), evaluating each section when held out.

### Lymphoid neighbourhood states redirect TLS maturation readouts

Two readouts were derived from the class logits: organization, (zETLS + zMTLS)/2 − zINFL, and maturity, zMTLS − zETLS. Across the five primary sections, organization separated TLS from infiltrate with a median spot AUC of 0.965, and maturity separated mature from early TLS objects with a mean AUC of 0.81 (Supplementary Fig. 3S1). In section LC3, a lymph node that was never a training target lay at the top of both readouts (median maturity: early TLS −2.00, mature TLS −1.38, germinal centre −0.27, lymph node 0.18; Fig. 3a,b). We then kept each observer’s H&E patch and centre gene list and replaced only its neighbour gene list with that of a real donor spot from the same section. The maturity reading moved systematically with the donor state (Fig. 3c). In LC3, early-TLS observers moved by −0.45 with infiltrate neighbourhoods, +0.21 with mature-TLS, +0.87 with germinal-centre and +1.22 with lymph-node neighbourhoods. Across the five sections, infiltrate, normal and tumour donors lowered the reading by 0.40–0.46, whereas mature-TLS donors raised it by 0.43, germinal-centre donors by 0.60 (two sections) and lymph-node donors by 1.22 (one section). Swapping the neighbourhood alone thus closed nearly half of the 0.96 difference between early and mature TLS. Conversely, across the five sections, mature-TLS observers dropped by 0.88 with infiltrate neighbourhoods, and infiltrate observers gained 0.72 in organization with mature-TLS neighbourhoods (Supplementary Fig. 3S2).

### Perturbation resolves opposing follicular B-cell and plasma-cell secretory programmes

To identify the neighbourhood states that move the reading, we correlated, within lymphoid donors, the change each donor caused with the measured whole-transcriptome expression of its neighbourhood, partialling out neighbourhood B-cell, T-cell and PTPRC expression. Co-expression clustering yielded two programmes (Fig. 3d). A follicular B-cell programme (FB; 20 genes, including NIBAN3, CD37, SPIB, BANK1, IRF8, CCR7, FCRL1 and CIITA) combined B-cell identity with MHC class II antigen presentation and raised the maturity reading. A plasma-cell secretory programme (PC; 20 genes, including IGHG1, IGKC, JCHAIN, XBP1, ERN1, SEC11C, DERL3 and SDC1) combined immunoglobulin chains, the IRE1–XBP1 unfolded-protein response and CD138, and lowered it. When both programmes were re-derived on four sections and tested on the fifth, FB was positive (partial ρ = 0.06 to 0.39) and PC negative (−0.24 to −0.49) in all 15 held-out tests (five sections × three observer–readout combinations; LC3: FB +0.19, PC −0.29; Fig. 3e; Supplementary Fig. 3S3). In measured neighbourhood expression, FB increased from infiltrate to early and mature TLS in four of five sections and, in LC3, further to germinal centres and the lymph node. VINTER therefore reads TLS maturation from broad neighbourhood programmes that separate follicular B-cell identity from plasma-cell secretion, rather than from germinal-centre transcripts, which are sparse on FFPE Visium (AICDA was detected in 2.5% of mature spots).

Editing the programmes directly changed the reading (Fig. 3f). In 34,697 observer–donor pairs (351 observers in five sections, up to 100 real lymphoid donors per section), we deleted programme genes from the donor’s neighbour list or inserted the donor’s three highest-ranked programme genes at the top. All four edits acted in the expected direction: deleting PC raised maturity (+0.36 and +0.37 for early- and mature-TLS observers), inserting PC lowered it (−0.27 and −0.26), deleting FB lowered it (−0.32 and −0.28) and inserting FB raised it (+0.32 and +0.30). Each edit shifted the reading in the same direction in all five sections and in 95–100% of observers. Because inserting three genes moved the reading in opposite directions for FB and PC, the effect reflects programme identity rather than the edit itself. Deleting PC from PC-high donors closed about 40% of their gap to PC-low donors, and inserting FB into FB-low donors closed about 44% of their gap to FB-high donors. Effect sizes varied between sections, from about ±1 in KC3 to within 0.1 in LC2 and LC5 (Supplementary Fig. 3S4).

### Spatial mapping reveals follicular interiors and secretory peripheries of mature TLS

Why would plasma-cell secretion, a product of mature TLS, lower the maturity reading? Mapping both programmes in tissue resolved this (Fig. 3g,h). In the mature TLS LC3_TLScc009, FB decreased from the core to the first outer ring (0.86, 0.59 and 0.29 in core, edge and ring 1), whereas PC increased outward (1.38, 1.77 and 1.54). PC was higher in ring 1 than in the core in all eight mature TLS of LC3 and in 17 of 17 mature TLS across the five sections (edge higher than core in 16 of 17), while FB was concentrated inside the TLS (edge higher than ring 1 in 11 of 17; Supplementary Fig. 3S5). This inside–outside separation characterized mature TLS: the median ring-1-minus-core PC difference was +0.83 in mature TLS (17 objects) and −0.06 in early TLS (6 objects with a core), and PC in ring 1 of mature TLS exceeded that of early TLS in all five sections. The model’s own maturity reading likewise decreased from core (1.34) to edge (−0.01) and ring 1 (−0.74). Plasma-cell secretion associated with mature TLS is therefore enriched at the TLS periphery, whereas the maturity that VINTER reads at spot scale arises from the follicular interior; neighbourhoods rich in PC accordingly pull an observer towards an early reading. This organization is consistent with B-cell maturation within TLS and the dissemination of plasma cells along fibroblast tracks into the tumour bed in renal cell carcinoma (45). VINTER resolves it from its maturity judgement through editable neighbourhood evidence.

### The follicular programme marks organized lymphoid nodules in an independent breast cancer cohort

The FB programme, defined on lung and kidney sections, also marked organized lymphoid nodules in the independent TNBC cohort (legacy spatial transcriptomics; 92 primary tumours) (18) (Fig. 3i,j). It shared 5 of 20 genes with the authors’ breast TLS signature (BLK, CCR7, CD37, FCMR and NIBAN3; hypergeometric P = 9.9 × 10⁻¹¹). FB was higher in nodules than in the first ring in 19 of 20 arrays with nodules, and higher in nodules than in ordinary immune infiltrate in 17 of the 18 patients with both. The plasma-cell periphery was not reproduced in this cohort (ring 1 higher than nodules in 6 of 20 arrays; Supplementary Fig. 3S6), and the patient-level survival associations of FB and PC are provided in Supplementary Fig. 3S7. These perturbations edit the gene-name lists supplied to the model and read its judgement; they are not tissue interventions, and TLS objects are connected regions of author labels.

### VINTER’s class probabilities reveal stromal immune heterogeneity and recover overlooked cancer in triple-negative breast cancer

Spatial annotation methods usually report a single label per spot, yet the tissue beneath a label is rarely uniform. Stroma in triple-negative breast cancer (TNBC) ranges from lymphocyte-dense to immune-silent, and stromal tumour-infiltrating lymphocytes (TILs) are among the best-established immune biomarkers of the disease. Because VINTER returns a probability for every tissue class, we asked whether these class probabilities retain this information. We applied the HER2ST-trained model, without any TNBC training, to primary tumours from 92 patients profiled by spatial transcriptomics with pixel-level pathology annotation (18).

### The class posterior resolves immune heterogeneity within uniformly labelled stroma

Within regions that VINTER assigned uniformly to connective tissue, the posterior probability of immune infiltrate varied over several orders of magnitude and followed the pathologists’ distinction between high- and low-TIL stroma (Fig. 4a). p(immune) distinguished high-from low-TIL stromal spots with a mean within-patient AUC of 0.87 (11 patients, all above 0.5) and tracked the continuous high-TIL fraction (mean Spearman’s ρ = 0.40; positive in 17 of 17 patients; Fig. 4b). The immune posterior is therefore a quantitative readout of stromal immune heterogeneity that the hard tissue label flattens.

### Posterior-defined hot stroma harbours a T-cell-inflamed transcriptional programme

We divided each patient’s stroma into posterior-hot, intermediate and cold tertiles and compared paired whole-transcriptome pseudobulks. Hot stroma showed 2,059 and cold stroma 1,261 up-regulated genes (FDR < 0.05; 92 patients; Fig. 4c). Among genes that VINTER never received as input, hot stroma was enriched for antigen processing (higher in 80 of 92 patients), IFN-γ-related genes including STAT1, CXCL9 and CXCL10 (80/92), the cytolytic effector PRF1 (62/92) and lymphoid-organising chemokines (66/92), whereas cold stroma was marked by lipid-handling and adipocyte genes such as PLIN4, GPD1, CD36 and LIPE (Fig. 4c,d). These features match the T-cell-inflamed microenvironment associated with response to immune checkpoint blockade. A zero-shot class posterior, read within a single hard label, thus captures a molecular axis of stromal immune activation that extends beyond the genes supplied to the model.

### Hot stroma separates into lymphoid-organised and IFN-dominated environments

Posterior-hot stroma was not uniform. We compared hot stroma lying next to author-annotated lymphoid nodules (within two spot spacings) with hot stroma far from them (more than four spacings), excluding from both groups any spot that contained nodule pixels. In the nine patients with both environments, hot stroma next to nodules was enriched for the lymphoid-organising chemokines CCL19, CCL21, CXCL13 and LAMP3 and for LTB (7 of 9 patients each), whereas hot stroma away from nodules carried a stronger IFN response (CXCL9, CXCL10 and STAT1; higher in 8 of 9 patients; Fig. 4e,f). When spots at nodule edges were retained, the organising signal was higher next to nodules in 11 of 12 patients (Supplementary Fig. 4S1). VINTER’s posterior therefore marks both environments as immune-hot, and spatial context separates lymphoid organization, reminiscent of the tertiary lymphoid structures analysed in Fig. 3, from IFN-dominated inflammation away from lymphoid aggregates.

### Posterior-guided review recovers nearly half of the cancer concealed by benign labels

The same principle applies to malignant tissue. In a representative tumour, 389 of 398 spots that pathologists annotated as cancer but VINTER labelled as non-tumour were assigned to breast glands, while their secondary cancer probability remained informative (Fig. 4g). Ranking non-tumour spots by this probability and reviewing 10% of each patient’s spots recovered on average 48% of such missed cancer across 53 patients, compared with 25% for random review at the same budget; at a 20% budget recovery rose to 75% (Fig. 4h). VINTER thus converts secondary class probabilities into an effective prioritization strategy for cancer that the hard label conceals.

### Cancer missed by hard labels is less proliferative, less epithelial and concentrated in mesenchymal TNBC

The cancer recovered by review was not a random subset. Compared with cancer that VINTER labelled correctly in the same 43 patients, and despite a similar pathologist-annotated cancer area (0.71 versus 0.74), recovered cancer showed lower proliferation programmes (E2F and G2M targets lower in 39 of 43 patients), weaker epithelial identity (35/43) and stronger stromal and immune transcripts (Fig. 4i). At the patient level, the fraction of high-cancer-area spots labelled non-tumour was highest in the mesenchymal molecular subtype and discriminated it from the other subtypes (AUC 0.76; 68 patients; Fig. 4j); the association persisted after adjustment for cancer burden, spot number and grade (P = 0.0005; 66 patients with complete covariates). Hard-label errors in TNBC are therefore biologically structured: the regions a single most-likely label misses are the least canonically epithelial, least proliferative parts of the tumour, and posterior-guided review recovers them.

Together, these analyses establish VINTER’s class probabilities as a biologically informative readout of tissue heterogeneity, in both the immune microenvironment and the tumour itself, and as a practical basis for prioritizing regions for pathological review in a new cohort without retraining.

### VINTER reads TLS state on Visium HD bins and Xenium cells without retraining

In Fig. 3, the follicular B (FB) programme raised VINTER’s maturity reading, the plasma-cell (PC) programme lowered it, and the PC programme was enriched around mature TLS. A Visium spot mixes several cells, however, so it could not show which cells carry these signals. Visium HD and Xenium resolve tissue at the scale of single bins and cells, but provide no TLS annotation to train on.

We therefore applied the Fig. 3 TLS model, without retraining, to colorectal cancer from three patients (20). Each patient was profiled by Visium HD and, on an adjacent section, by Xenium. Each 8-µm bin or cell was a centre unit, read with its real spatial neighbours and the surrounding H&E (Methods).

### VINTER identifies TLS on both platforms

VINTER’s TLS probability marked B-cell-rich tissue on both platforms (Fig. 5a,b): on Visium HD, the authors’ RCTD B-cell bins, with an AUC of 0.99; on Xenium, B cells, with an AUC of 0.93. Detection held in a second patient (Supplementary Fig. 5S1b). In a patient with almost no B cells, VINTER found no TLS structure. On the adjacent sections, the two platforms delineated the same TLS (Cohen’s κ = 0.81; Supplementary Fig. 5S1a).

### Maturity follows the follicular programme

Within TLS, the maturity reading was highest over the follicles and followed the measured FB programme (Fig. 5c). On Xenium, B cells filled the TLS interior, with T cells interspersed (Fig. 5d).

### Plasma cells gather at TLS margins

On both platforms, plasma cells were sparse inside TLS and enriched at their margins (Fig. 5e–g). Visium HD resolves isotypes, which the Xenium panel cannot: plasma cells at the TLS margin were mainly IgG⁺, and plasma cells in the mucosa and at the tumour border were mainly IgA⁺ (Fig. 5e; Supplementary Fig. 5S1d).

### Cell-level readings follow the **Fig.** 3 programmes

Every cell type read as more mature inside TLS than outside, and at the same distance from the edge plasma cells read lower than B cells (Fig. 5h).

We then replaced only a cell’s own gene list and kept its H&E field and neighbourhood (Fig. 5i). A follicular B cell given a plasma cell’s list read as less mature, a rim plasma cell given a B cell’s list read as more mature, and exchanging lists between B cells had no detectable effect. The shifts followed the direction found in Fig. 3. They were modest, and they left the TLS calls essentially unchanged.

Together, these results place the observation of Fig. 3 on cells. Plasma-cell transcripts lower the maturity reading, and in tissue, plasma cells sit at the margins of TLS, where they are mainly IgG⁺. This agrees with IgG⁺ plasma cells reported around TLS in colorectal liver metastases (46) and with TLS as a source of IgG-producing plasma cells in renal cancer (45).

## Discussion

VINTER is a generative vision–language model that reads histology, measured gene expression and spatial neighbourhood as a single sequence. Visual–omics and spatial foundation models have shown the value of linking molecular measurements with tissue morphology and neighbourhood structure (4–7). VINTER develops this direction by making each tissue decision a joint reading of all three sources and by keeping the evidence behind that decision readable and editable. This design annotated cortical layers more accurately than spatial-domain methods and a spatial foundation model, and transferred to an independent breast cancer cohort without target labels (Fig. 2). It resolved the neighbourhood programmes behind a judgement of tertiary lymphoid structure (TLS) maturation (Fig. 3), exposed immune and malignant heterogeneity concealed by single tissue labels (Fig. 4), and read Visium HD 8-µm bins and Xenium single cells without retraining (Fig. 5).

Representing gene profiles as text is what turns annotation into interrogation. Gene names preserve recognizable molecular identities, their order conveys relative expression, and a separate neighbourhood list places each spot in its local tissue context; text-based gene representations have proved effective in related work (47). Because the neighbourhood enters as its own list, it can be edited, or replaced by the neighbourhood of another real spot, while the image and the spot’s own genes are held fixed. In silico perturbation with single-cell foundation models nominates genes that shift the state of a cell (14,15); VINTER instead asks which neighbourhood evidence shifts a tissue-level judgement. For TLS, this question separated a follicular B-cell programme that raised the maturity reading from a plasma-cell secretory programme that lowered it. Mapping both programmes in tissue resolved the apparent paradox: plasma-cell secretion, a product of mature TLS, was concentrated at the periphery of the structure, whereas the maturity read at spot scale arose from the follicular interior. This organization agrees with the view of TLS as sites that generate and disseminate antibody-producing plasma cells (19,45), and the follicular programme defined in lung and kidney cancer also marked organized lymphoid nodules in breast cancer. Neighbourhood perturbation therefore moves from explaining a model decision to nominating spatial hypotheses that can be tested in independent tissue.

An informative annotation must also retain what a single label discards. VINTER returns the probability of every candidate class, and in triple-negative breast cancer (18) these posteriors, read within regions that shared one label, recovered structure that pathologists recognize. Within uniformly labelled stroma, the immune posterior followed the pathologists’ distinction between high- and low-TIL stroma, an established prognostic feature of the disease (11). Posterior-hot stroma carried T-cell-inflamed and interferon-response programmes among genes that the model never received, and separated lymphoid-organized environments next to nodules from interferon-dominated inflammation elsewhere. In benign-labelled regions, secondary cancer probabilities directed review to nearly half of the concealed cancer at a 10% budget; the recovered cancer was less proliferative and less epithelial, and hard-label misses were concentrated in mesenchymal tumours. Class posteriors thus serve as a representation of tissue heterogeneity in their own right, complementing approaches that carry prediction uncertainty into downstream spatial analyses (12), and they offer a practical way to prioritize pathological review in a new cohort without retraining.

The same design makes VINTER applicable across tissues, platforms and resolutions. In this study, one architecture and interface was applied to human cortex, breast, lung, kidney and colorectal cancers; read legacy spatial transcriptomics with 100-µm spots, Visium, Visium HD and Xenium; and served three tissue vocabularies (cortical layers, breast tissue classes and TLS stages) through the same class-token head. Two properties underlie this reach. Genes enter as a ranked list of names computed from the library-normalized profile of each unit, so the input keeps the same form whatever the capture depth or unit size, and the image branch builds on a general-purpose pathology foundation model (17). Accordingly, the TLS model trained on 55-µm Visium spots from lung and kidney cancer read colorectal cancer Visium HD bins and Xenium cells without retraining (20). It identified TLS in close agreement with B-cell annotations, reproduced the follicular and plasma-cell relationships learned on Visium, resolved two follicles within a single TLS at 8 µm, and located IgG⁺ plasma cells at TLS margins, whereas IgA⁺ plasma cells predominated in the mucosa and at the tumour border. These results closed the loop opened by perturbation: the plasma-cell programme that neighbourhood editing nominated, and that mapped to the periphery of mature TLS on 55-µm spots, appeared in 8-µm plasma-cell bins as a class-switched IgG⁺ niche rimming VINTER-defined TLS. Resolution thus becomes a choice made at inference rather than a property fixed at training, so that models trained on abundant annotated spot-level data can be carried to higher-resolution platforms for which annotations do not yet exist.

Three directions would extend this framework. First, the downstream analyses rest on published annotations in selected cohorts; TLS stages in particular followed the labels of the original study, and stain-based staging in larger cohorts would sharpen the maturity reading and allow it to be related to clinical outcome. Second, VINTER reads each neighbourhood as one aggregated gene list; representing individual neighbours and their arrangement would let the model resolve layered structures, such as the follicle, mantle zone and T zone of a TLS, directly from its input. Third, input edits reveal the evidence that VINTER uses, and the programmes and niches they nominate, including the IgG⁺ plasma-cell margin of TLS, now warrant protein-level and functional validation.

Future work could extend pretraining to imaging-based platforms with single-cell resolution and use the class-token interface to add tissue vocabularies from a few labelled examples. Pairing posterior-guided review with prospective pathology, and model-derived programmes with targeted measurements, would bring these readouts into diagnostic and experimental use.

### Conclusions

VINTER establishes a generative vision–language framework in which histology, measured gene expression and spatial context are read together, and the evidence behind each tissue judgement can be inspected, edited and carried across platforms and resolutions. This framework turns spatial annotation into an interrogable interpretation of tissue organization, from annotated cohorts to new high-resolution technologies.

## Data Availability Statements

This study analysed previously published spatial transcriptomics datasets. DLPFC data are available through the spatialLIBD resource; HER2ST data through https://zenodo.org/records/3957257; TNBC data through https://zenodo.org/records/8135721; the kidney and lung cancer Visium sections with TLS annotations through https://zenodo.org/records/14620362; and the colorectal cancer Visium HD and Xenium data through https://www.10xgenomics.com/products/visium-hd-spatial-gene-expression/dataset-human-crc (raw sequencing data, GEO accession GSE280318).

## Supporting information

Supplementary Information for VINTER

## Acknowledgements

This research was conducted using the Supermicro ARS-111GL-DNHR-LCC and FUJITSU Server PRIMERGY CX2550 M7 (Miyabi) at Joint Center for Advanced High Performance Computing (JCAHPC). Computational resources were also provided by the supercomputer system SHIROKANE at the Human Genome Center, Institute of Medical Science, University of Tokyo.

## Author Contributions

Conceptualization, J.W., Y.Y. and K.N.; methodology, J.W., W.X. and Y.Y.; software, J.W.; formal analysis, J.W., W.X. and Yizhou Zhuang; investigation, J.W.; data curation, J.W.; validation, W.X., Yizhou Zhuang and Yawen Zhang; visualization, J.W.; writing original manuscript, J.W.; review and editing, J.W., W.X., M.L., Y.Y. and K.N.; supervision, Y.Y. and K.N.

## Declarations

### Ethics Statement

This study reanalysed published, de-identified spatial transcriptomics datasets. No new human participants were recruited and no new tissue specimens were collected. Ethics approval and consent procedures for the original studies are described in the corresponding publications.

### Consent for publication

Not applicable.

### Generative AI statement

The authors declared that Generative AI was not used in the creation of this manuscript.

## Abbreviations

ARI: adjusted Rand index;
AUC: area under the receiver operating characteristic curve;
CI: confidence interval;
DLPFC: dorsolateral prefrontal cortex;
ETLS: early tertiary lymphoid structure;
FB: follicular B-cell programme;
FDC: follicular dendritic cell;
FDR: false discovery rate;
FFPE: formalin-fixed paraffin-embedded;
GC: germinal centre;
H&E: haematoxylin and eosin;
HER2: human epidermal growth factor receptor 2;
IFN: interferon;
INFL: immune infiltrate;
LN: lymph node;
LoRA: low-rank adaptation;
MTLS: mature tertiary lymphoid structure;
NMI: normalized mutual information;
PC: plasma-cell secretory programme;
RCTD: robust cell type decomposition;
ST: spatial transcriptomics;
Tfh: T follicular helper;
TIL: tumour-infiltrating lymphocyte;
TLS: tertiary lymphoid structure;
TNBC: triple-negative breast cancer;
VINTER: Vision–language Interrogation of Neighborhood Transcriptomic Evidence and Reasoning;
WM: white matter.

