## Supplementary Information for VINTER for "VINTER: a generative vision–language model for annotating and interrogating spatial tissues"

Supplementary Figures and Tables are numbered by the main figure they support (for example, Supplementary Fig. 2S1 accompanies Fig. 2).

### Supplementary Figures

**Supplementary Figure 2S1 | Balanced accuracy, ARI and NMI.**

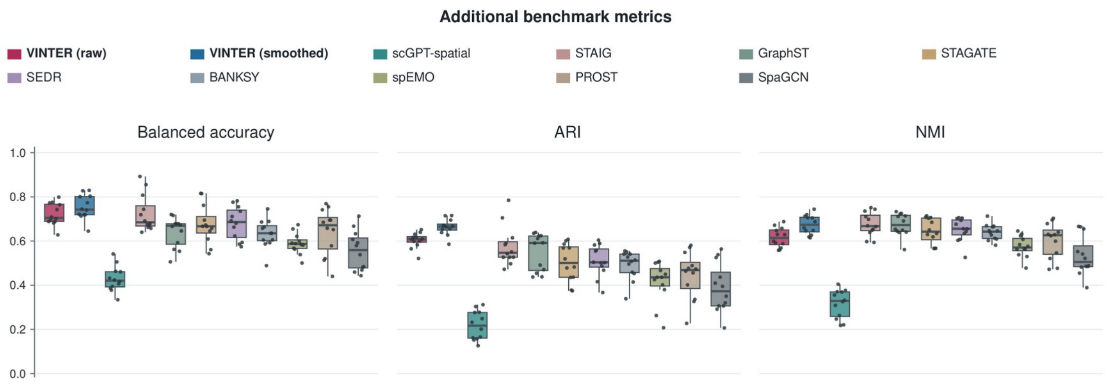


Twelve section-level values per output, following the main benchmark's EVAL scope, method order and box definitions.

**Supplementary Figure 2S2 | Layer-specific F1.**

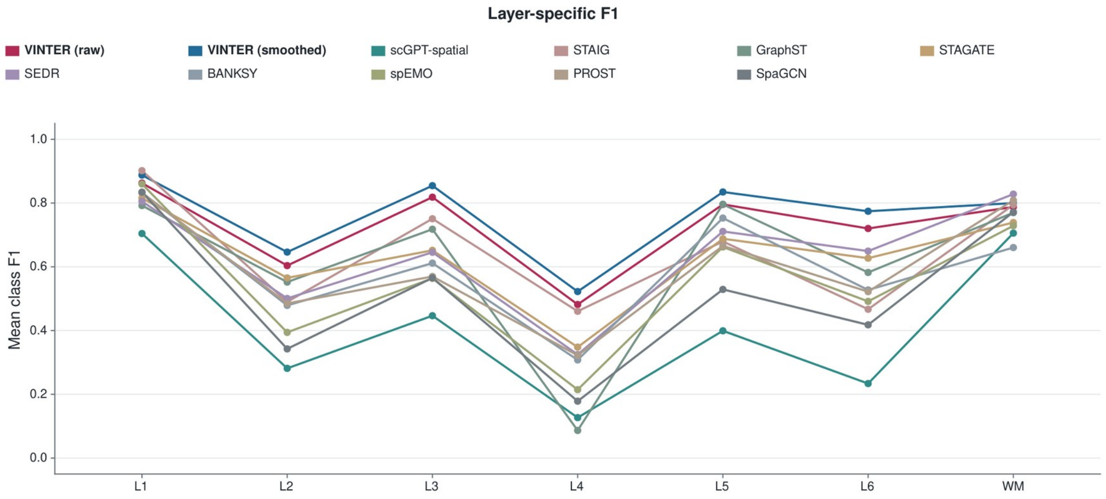


Measured class F1 averaged with equal section weight among sections with positive reference support for each class: eight sections for L1/L2 and twelve for L3–L6/WM. BANKSY uses the fixed displayed run rather than resolution-grid medians.

**Supplementary Figure 2S3 | Multimodal complementarity.**

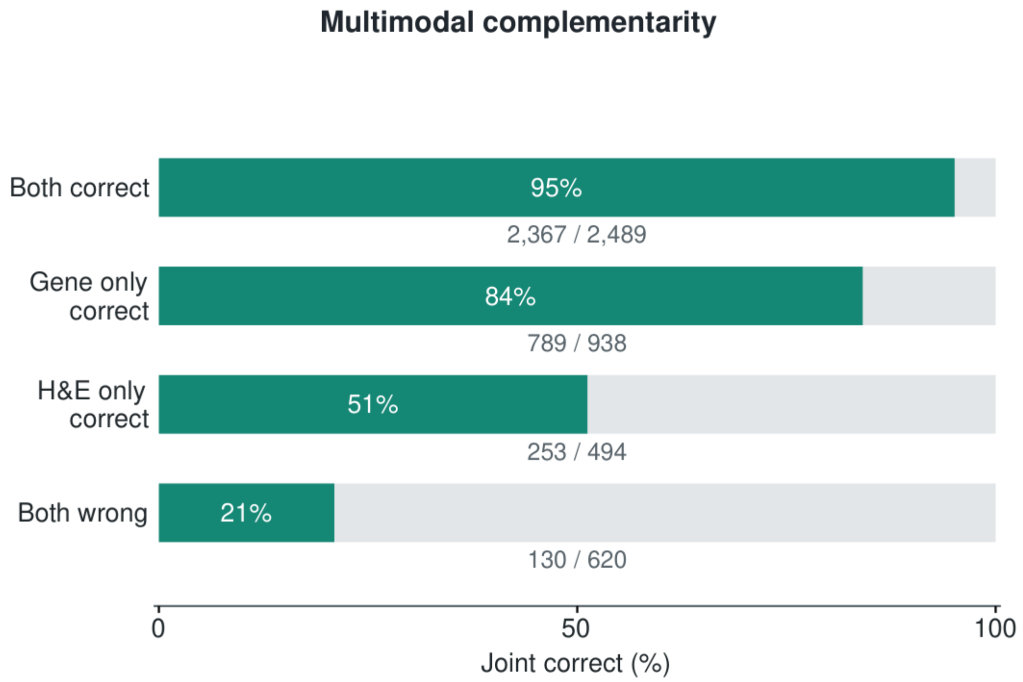


All 4,541 EVAL spots from sections 151675 and 151676 are grouped by the correctness of gene-only and H&E-only predictions. Bars report raw-joint-model correctness within each group; gray remainders are errors. All four groups and their exact denominators are retained.

**Supplementary Figure 2S4 | Input-edit sensitivity distribution.**

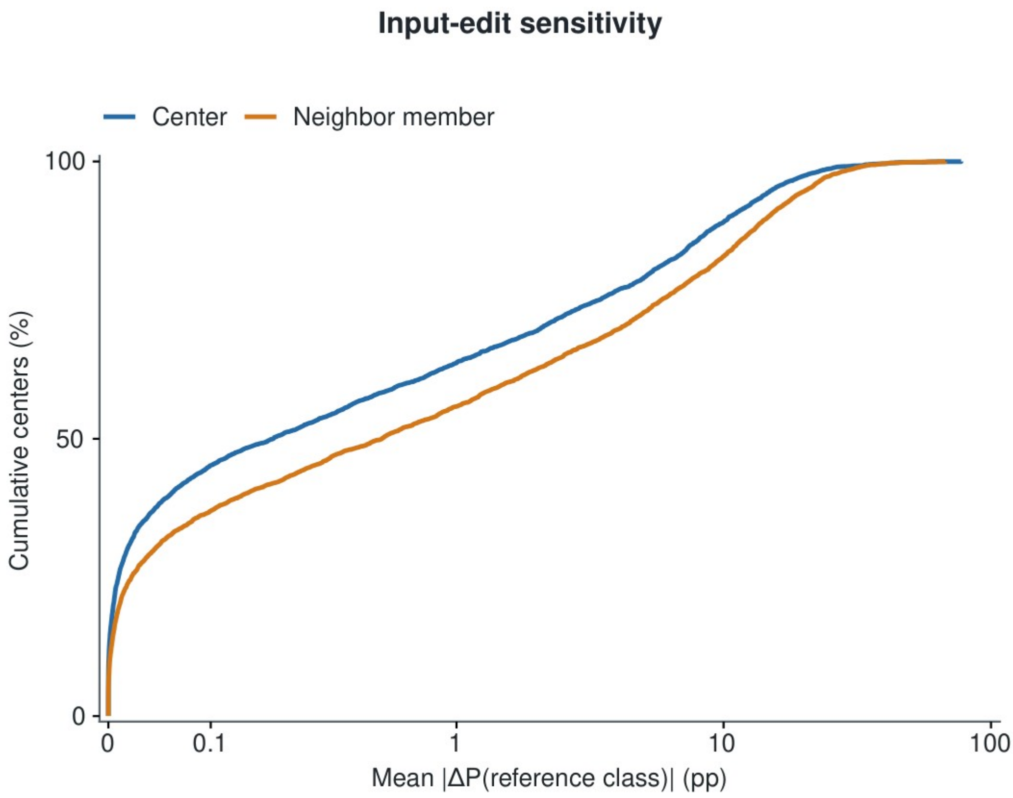


For each of 4,541 EVAL centers, the mean absolute change in reference-class probability is calculated over expression-zero-and-rerank events: twenty center genes, or ten pooled-neighborhood genes edited individually in each of six neighbor members. No-op events remain in the denominator. Curves are empirical cumulative distributions with a pseudo-log x axis. This is an aggregate sensitivity diagnostic. Its expression-zero/repooling operator differs from the direct ranked-name edits shown in Fig. 2f.

**Supplementary Figure 2S5 | VINTER adaptation.**

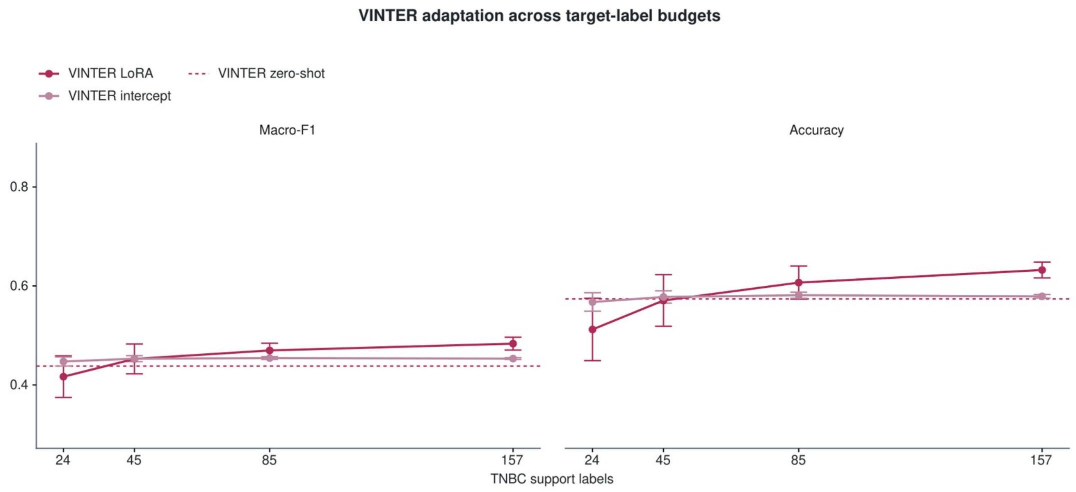


LoRA and intercept adaptation are shown at 24, 45, 85 and 157 labels, with five support draws at each budget. Points are equally weighted test-patient means averaged over draws; error bars are one standard deviation across support draws. Dashed lines are zero-shot references. All support labels originate from eighteen adaptation patients and are disjoint from the 56 test patients. The protocol also contains eighteen validation patients; LoRA uses 128 fixed update steps without validation/test-based early stopping. These are within-model adaptation results.

**Supplementary Figure 2S6 | Exact rank-edit responses and complete section summary.**

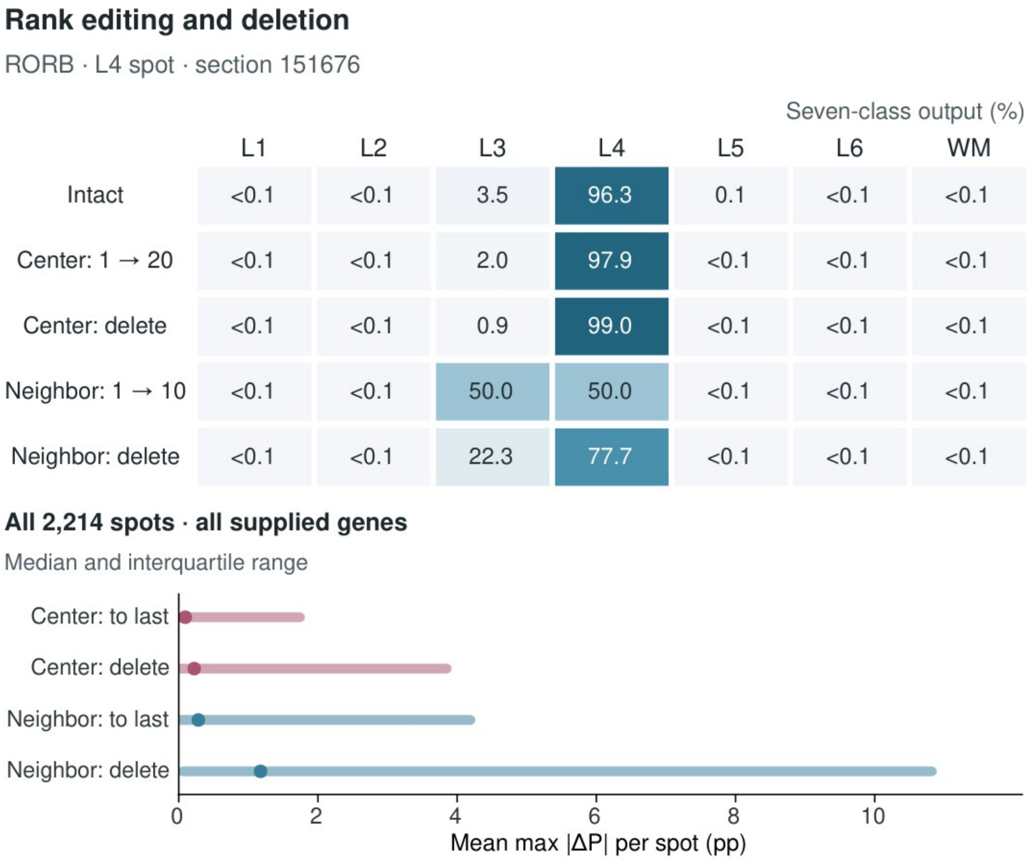


The upper matrix reports all seven output probabilities for the intact input and four RORB edits shown in Fig. 2f. Numbers are percentages; values below 0.1% are marked accordingly. The lower plot summarizes all 2,214 canonical EVAL spots in section 151676. For each spot and operation, the maximum absolute change over the seven output classes is computed per supplied gene, then averaged across all twenty center or ten pooled-neighbor genes. Points are medians and ranges are interquartile ranges across spots. All 132,840 measured events are included, including zero responses and unchanged prompts when an already-last gene is moved to the last position. The two edit universes differ in gene sets and list lengths and do not establish modality importance.

**Supplementary Figure 2S7 | Complete six-channel probability atlas.**

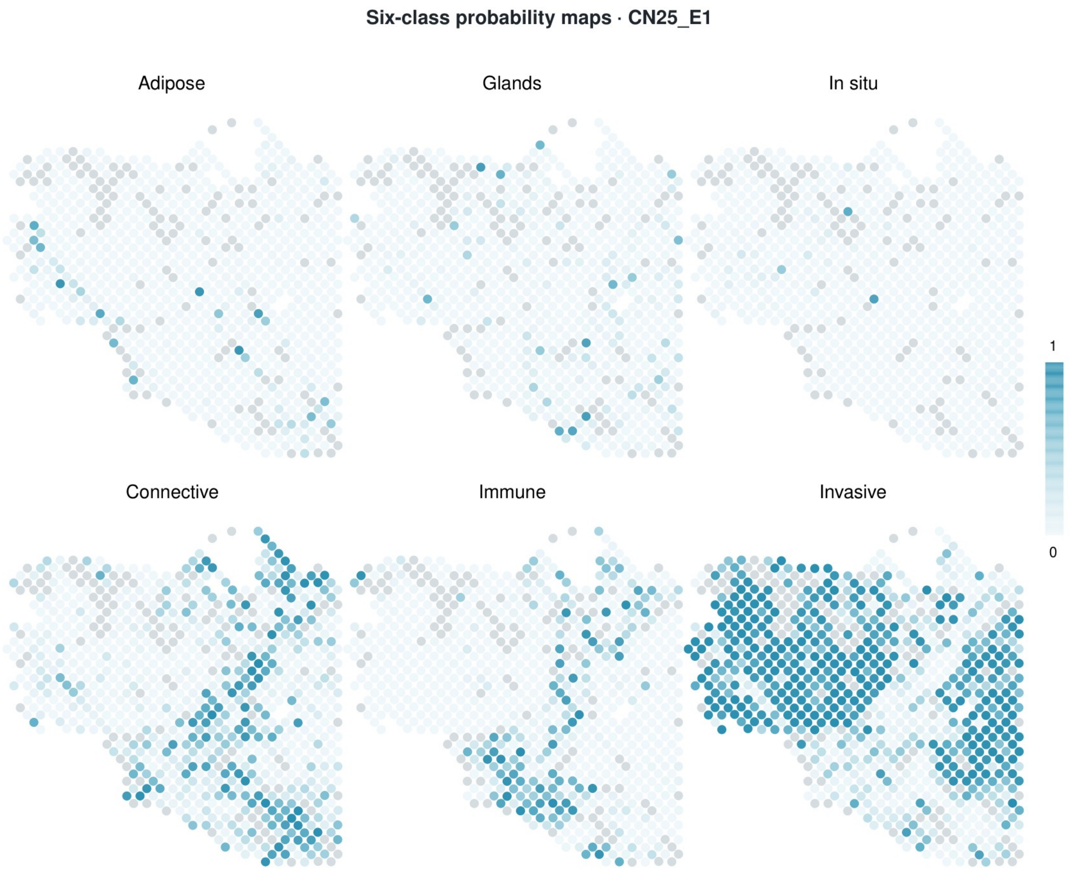


All six raw restricted-softmax channels for CN25_E1, sharing one 0–1 color scale and exactly the same coordinate window, unmodified spot centers and disk sizes as Fig. 2c,e. All 812 measured positions are displayed, including the 116 excluded positions in gray. No spatial interpolation or additional spots are introduced.

**Supplementary Figure 2S8 | Exact example vectors and confidence-based retention.**

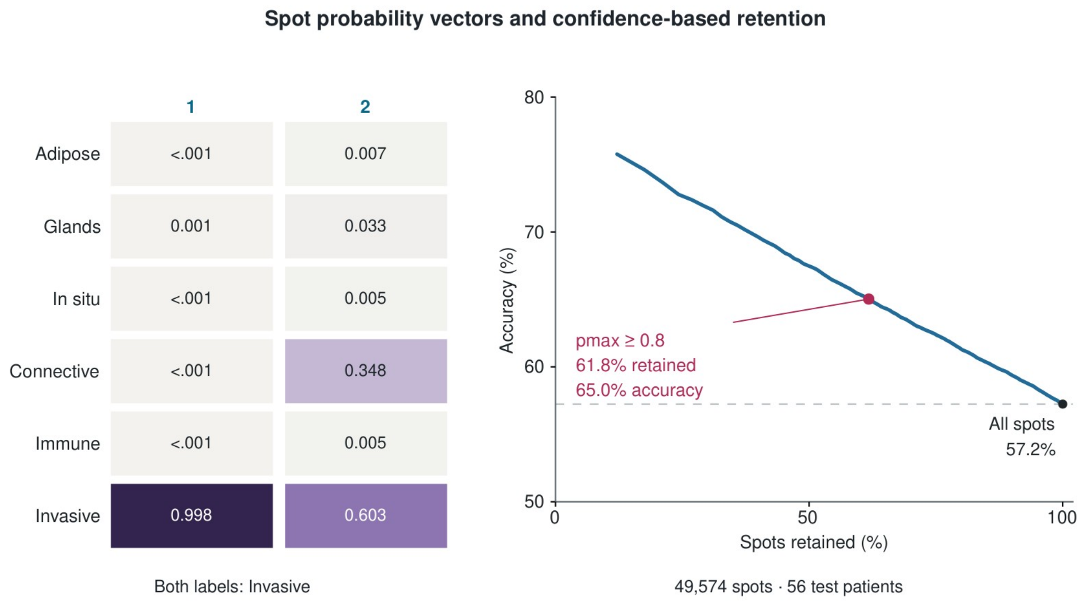


Left: full six-class vectors for spots 1 and 2 in Fig. 2e, shown in matching numeric order and rounded to three decimals; values below 0.001 are marked. Both have Invasive argmax. Right: pooled retained-spot accuracy against the fraction of eligible spots retained across the fixed 56-patient test cohort. Thresholds span 0–0.995 in steps of 0.005. At the illustrative threshold of 0.8, 30,614 of 49,574 spots are retained (61.8%), with 19,907 correct predictions (65.0%), versus 57.2% over all eligible spots. The source-domain temperature is not applied. This retrospective accuracy–coverage trade-off is descriptive, not target-domain calibration or a validated clinical decision rule.

**Supplementary Figure 3S1 | Staging performance of the five-class TLS model.**

*Figure in preparation.*

Complete organization and maturity metrics for all sections, comparison with the four-class model, and the model retrained without germinal-centre, FDC and Tfh genes (object maturity AUC 0.712 versus 0.716 for the full model).

**Supplementary Figure 3S2 | Complete real-neighbourhood swap matrix.**

*Figure in preparation.*

All observer–donor class combinations for both readouts in the five primary sections, and the prespecified programme scan.

**Supplementary Figure 3S3 | Follicular B and plasma-cell programme genes.**

*Figure in preparation.*

All 40 programme genes, per-gene partial correlations, co-expression clustering and the complete leave-one-section-out table.

**Supplementary Figure 3S4 | Bidirectional programme editing in all sections.**

*Figure in preparation.*

Deletion and insertion effects of FB and PC for every observer group in all five primary sections, and gap closure.

**Supplementary Figure 3S5 | Programme profiles of all mature TLS.**

*Figure in preparation.*

Core, edge and ring profiles for all 17 mature TLS and programme maps for all sections.

**Supplementary Figure 3S6 | TNBC nodules, rings and ordinary infiltrate.**

*Figure in preparation.*

FB and PC scores in nodule, ring 1, ring 2 and ordinary immune infiltrate for all primary TNBC arrays, including the non-reproduced PC periphery (ring 1 above nodule in 6 of 20 arrays).

**Supplementary Figure 3S7 | Survival associations of the TLS programmes in TNBC.**

*Figure in preparation.*

Patient-level overall-survival associations of the FB and PC programmes.

**Supplementary Figure 4S1 | Hot stroma next to lymphoid nodules when nodule-edge spots are retained.**

*Figure in preparation.*

The next-to versus away comparison of Fig. 4e,f with spots containing nodule pixels retained (12 patients; organizing chemokines higher next to nodules in 11 of 12).

**Supplementary Figure 5S1 | Cross-platform agreement, a second patient and further readouts for Fig. 5.**

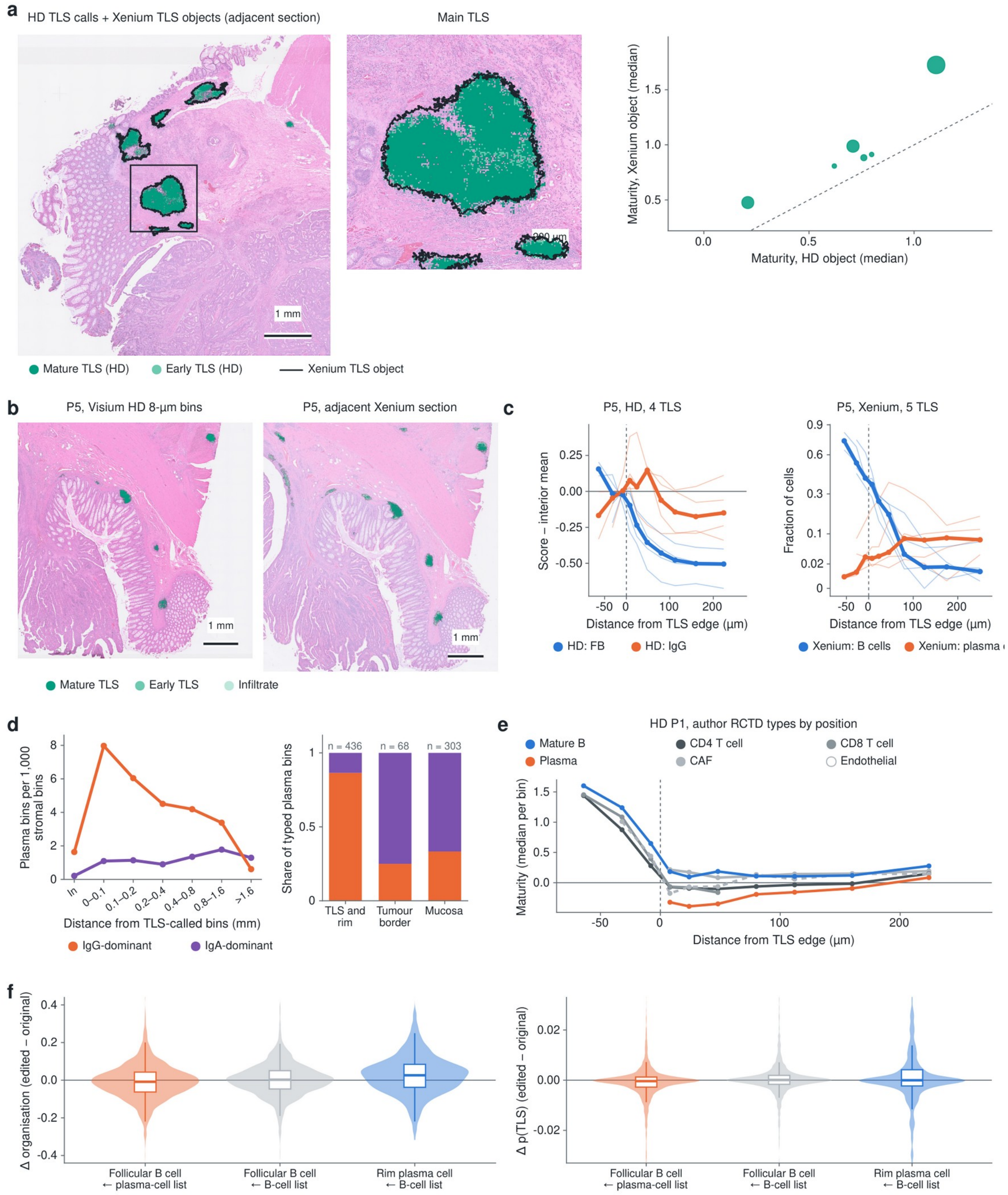


**a**, Cross-platform agreement in P1. VINTER TLS calls on Visium HD, overlaid with the TLS objects found on the adjacent Xenium section after image registration. Right: maturity of matched TLS objects on the two platforms.

**b**, Second patient, P5: VINTER lymphoid calls on Visium HD bins and on Xenium cells.
**c**, P5 profiles by distance from the TLS edge, as in Fig. 5g.
**d**, P1, Visium HD. Left: density of IgG- and IgA-dominant plasma bins by distance from TLS. Right: isotype composition of plasma bins in TLS and their rim, at the tumour border and in the mucosa.
**e**, P1, Visium HD: maturity of author-annotated cell types by distance from the TLS edge.
**f**, Centre-gene-list replacement as in Fig. 5i: change in the organization readout and in TLS probability.

### Supplementary Tables

**Supplementary Table 5S1 | Marker genes used to type Xenium cells.**

| Cell type | Marker genes |
|---|---|
| B cells | *MS4A1, CD79A, CD79B, PAX5, BANK1, FCRL1, FCRLA, SPIB, TCL1A* |
| Germinal-centre B cells (merged into B cells) | *RGS13, LRMP* |
| T cells | *CD3D, CD3E, CD3G, TRAC, TRBC1, TRBC2, CD2, CD5, CD6, IL7R, CD8A, CD8B* |
| Plasma cells | *MZB1, TNFRSF17, DERL3, FKBP11, SEC11C, PRDX4, IGHG3* |
| Myeloid cells | *CD68, CD163, C1QA, C1QB, C1QC, CD14, FCN1, MRC1, STAB1, ITGAX* |
| Fibroblasts | *PDGFRA, FAP, CXCL12, MFAP5, GREM1, COL11A1, CTHRC1, SOD3, RSPO3* |
| Smooth muscle cells | *ACTA2, DES, TAGLN, MYH14* |
| Endothelial cells | *PECAM1, VWF, CDH5, PLVAP, AQP1* |
| Epithelial cells | *CEACAM5, CEACAM6, PIGR, OLFM4, FABP1, LGR5, CA2, GUCA2A, SLC26A3, KRT80, CLCA1, REG4, TFF1* |
| Mast cells | *CPA3, MS4A2, KIT, HDC* |

All marker genes are on the 422-gene Xenium panel used for patients P1, P2 and P5. Each cell took the type with the highest marker score (summed marker counts divided by the square root of the number of markers); cells with too few marker counts were left unassigned. Germinal-centre B-cell markers were merged into B cells.
